# Cryo-EM structure of the bacterial effector protein SipA bound to F-actin reveals a unique mechanism for filament stabilization

**DOI:** 10.1101/2023.12.21.572903

**Authors:** Emily Z. Guo, Steven Z. Chou, Maria Lara-Tejero, Jorge E. Galán

**Affiliations:** Department of Microbial Pathogenesis, Yale University School of Medicine, New Haven, CT 06536; Department of Molecular Biology and Biophysics, UConn Health, Farmington, CT 06030

**Keywords:** bacterial pathogenesis, *Salmonella* pathogenesis, actin dynamics, type III protein secretion, host-pathogen interactions

## Abstract

The bacterial pathogen *Salmonella* spp. modulates cellular processes by delivering effector proteins through its type III secretion systems. Among these effectors, SipA facilitates bacterial invasion and promotes intestinal inflammation. The mechanisms by which this effector carries out these functions are incompletely understood, although SipA’s ability to modulate actin dynamics is central to some of these activities. Here, we report the cryo-EM structure of SipA bound to filamentous actin. The structure shows that this effector stabilizes actin filaments through unique interactions of its carboxy-terminal domain with four actin subunits. Furthermore, our structure-function studies revealed that SipA’s actin-binding activity is independent of its ability to stimulate intestinal inflammation. Overall, these studies illuminate critical aspects of *Salmonella* pathogenesis and provide unique insight into the mechanisms by which a bacterial effector modulates actin dynamics.

## MAIN

*Salmonella enterica* serovars can cause a range of human illnesses, from self-limiting gastroenteritis (i.e. food poisoning) to systemic, life-threatening infections (i.e. typhoid fever), resulting in an estimated 200,000 annual deaths worldwide (1–8). Central to *Salmonella* pathogenesis is their ability to invade intestinal epithelial cells and cause intestinal inflammation (4, 5, 7, 8). Epithelial cell invasion is orchestrated by the combined activity of several effectors of a type III protein secretion system (T3SS) encoded within its pathogenicity island 1 (SPI-1) (9–12). These effectors trigger cellular responses that result in the reorganization of the actin cytoskeleton, leading to the formation of large lamellipodia-like structures or membrane ruffles that internalize the bacteria into a membrane-bound compartment, the *Salmonella*-containing vacuole.

Three of these effector proteins, SopE, SopE2, and SopB, play a central role in the internalization process by stimulating Rho-family GTPases, which initiate downstream signaling leading to the WASH/WAVE-dependent activation of endogenous actin nucleators, such as the Arp2/3 complex (13–18). SopE, and its close homolog SopE2, directly target the Rho-family GTPases Rac1 and Cdc42, acting as potent exchange factors (13, 19). In contrast, SopB activates endogenous cellular exchange factors by fluxing phosphoinositides through its phosphoinositide-phosphatase activity (15).

Two other effectors, SipC and SipA, contribute to bacterial invasion by directly targeting actin (20, 21). Besides its role in T3SS-mediated protein translocation (22), SipC can nucleate and bundle actin filaments (21). A SipC mutant defective in actin-binding but proficient in T3SS-mediated protein translocation, showed a relatively small but significant defect in bacterial internalization (23), indicating an accessory but important role in bacterial cell entry. SipA plays a complex role in *Salmonella* pathogenesis. Through its carboxy-terminal actin-binding domain, SipA contributes to bacterial entry by reducing the critical concentration of actin required for its polymerization (20, 24). Furthermore, SipA stabilizes actin filaments and prevents the depolymerizing action of F-actin-severing proteins such as ADF/Cofilin and gelsolin (25). Consequently, the absence of SipA results in a reduced size of the bacterial-stimulated actin ruffles and a decrease in epithelial cell invasion (20, 25). In addition, through its amino-terminal domain, SipA modulates the intestinal inflammatory response to *Salmonella* by mechanisms that are not yet fully understood (26, 27). However, it remains unclear whether the actin-binding activity of SipA is related to this additional function since, despite previous efforts (28), the lack of high-resolution structural information on the SipA-actin interface has hampered the construction of SipA mutants that are essential to clarify this issue.

Similarly, due to the lack of structural information, the mechanisms by which SipA modulates actin dynamics are poorly understood. Although the crystal structure of the actin-binding domain of SipA in isolation is available (29), its atomic structure bound to F-actin is not. Previous low-resolution negative stained cryo-EM structures led to the proposal that SipA “staples” actin filaments through the deployment of two non-globular arms that reach out to opposing actin strands, presumably engaging actin in a similar manner to nebulin (29, 30). However, the lack of high-resolution structures has prevented the rigorous testing of such a model. Furthermore, machine learning–based approaches, such as that afforded by Alpha Fold (31, 32), are inadequate to predict the structure of SipA bound to F-actin. In this study, we report the cryo-EM structure of SipA bound to F-actin at 3.6 Å resolution. Our structure, validated by functional studies, clarifies the function of SipA, reveals a unique mechanism of F-actin stabilization by an actin-binding protein, and provides unique insight into the mechanisms of *Salmonella* invasion.

## Results

### SipA binding changes the symmetry of the decorated filaments

The *S.* Typhimurium effector protein SipA is 685 amino acids in length and is composed of at least three domains: an N-terminal chaperone-binding domain, a central ∼200 amino acid domain modeled by AlphaFold as long unstructured loops, and a C-terminal actin-binding globular domain (Supplementary Fig. 1). We first attempted to purify full-length SipA using different tags and purification conditions. However, consistent with previous reports (29), all constructs containing the central region were insoluble or unstable. Therefore, to study the actin-stabilizing function of SipA, we concentrated our efforts on its C-terminal domain (amino acids 447 to 685), which in previous studies was found to contain all its actin-modulating activities (20, 29). Consistent with previous reports (20, 29), we found that purified SipA_447-685_ interacts with skeletal muscle actin with high affinity (0.54 µM) as determined by a co-sedimentation assay (Supplementary Fig. 2). Also, consistent with previous observations (20), negatively stained actin filaments polymerized in the presence of SipA observed under the electron microscope showed a thicker and straighter appearance than filaments polymerized in its absence (Supplementary Fig. 3).

To determine the structure of the SipA-actin complex, purified G-actin was first polymerized to form filaments, mixed with purified SipA_447-685,_ and examined by single particle cryo-EM. We collected 4,740 movies, of which 2,370 were used for manual filament picking (see Methods). In total, 278,740 filament segments were boxed out for further analysis. The helical parameters of SipA_447-685_-decorated actin filaments were determined by helical indexing of the averaged power spectrum. The rise and translation of actin subunits along the helix of bare actin filaments is about 27.4 Å. Notably, the rise of SipA-decorated actin filaments is 109.9 Å, roughly four times longer than that of bare actin filaments. The rise line (the first layer line that crosses the meridian), although weak, is visible in the averaged power spectrum (Supplementary Fig. 4). By applying helical symmetry in the EM reconstruction, we obtained a map with extra densities (colored in magenta and yellow in Fig. 1) alongside the grooves between the two strands of the actin filaments (colored light green and dark green in Fig. 1), which were not present in actin filaments alone and therefore assumed to correspond to SipA (Fig. 1). Notably, in 2D class averages, the SipA densities were much weaker than that of the actin filaments and displayed low resolution and poor map connectivity in initial 3D reconstructions (Supplementary Fig. 5). These findings indicate a sub-stoichiometric presence of SipA on the actin filaments. We processed the particles with SipA binding at both grooves and generated a final map with a global resolution of 3.6 Å determined by the Fourier shell correlation (FSC) at 0.143 cutoff (Supplementary Fig. 6). To improve the map connectivity and interpretability, DeepEMhancer (33) was applied to post-process the refined 3D map. Local resolution analysis revealed that the cryo-EM map for the actin filament exhibits a higher resolution of approximately 3.5 Å, whereas the SipA map displays a lower resolution, ranging from 3.5 Å to 5.5 Å (Supplementary Fig. 7). This map showed densities corresponding to the side chains of most actin residues, as well as the bound nucleotide and the associated Mg2+ in polymerized actin (Supplementary Fig. 8).

**Figure 1.**
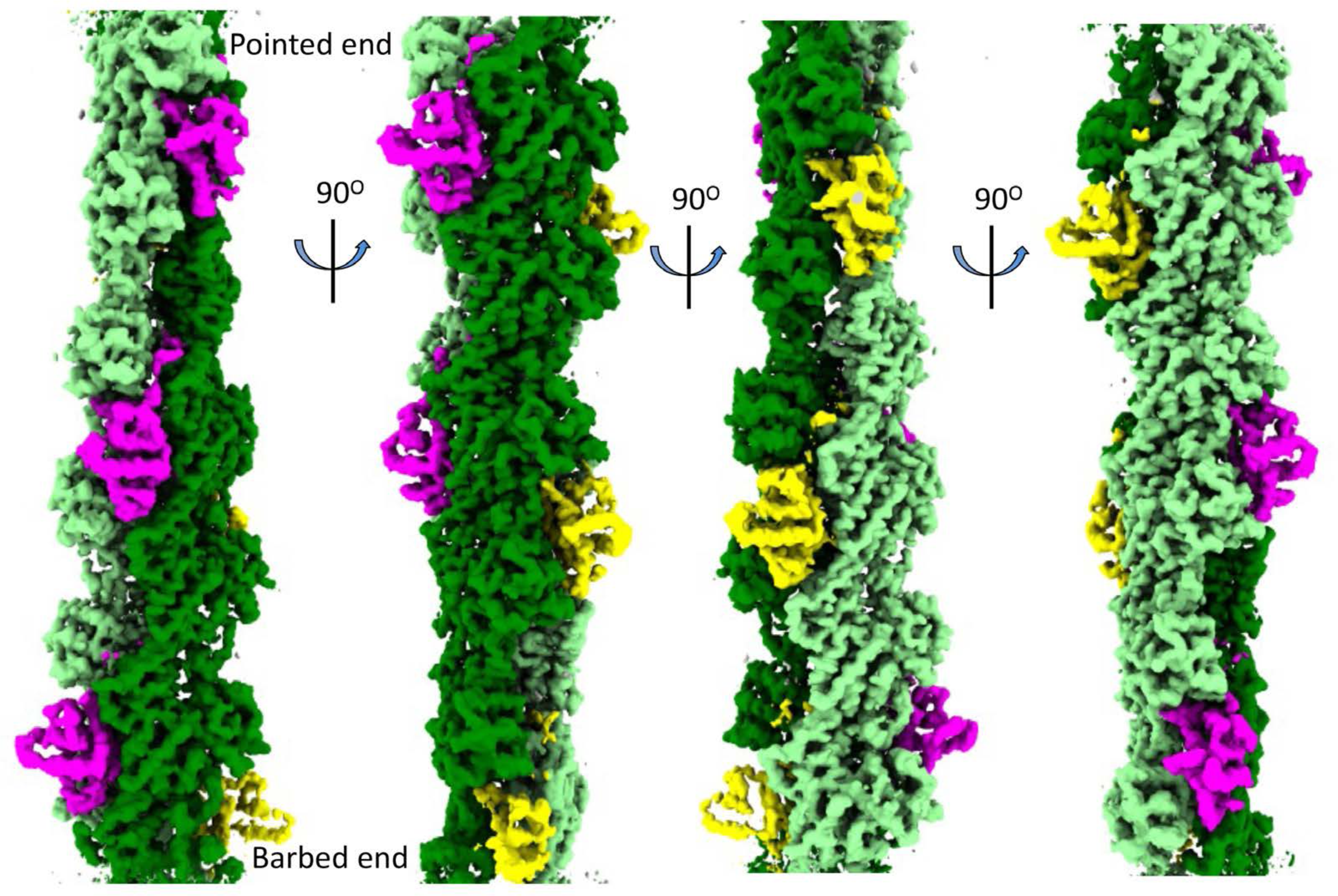
Overall structure of SipA_447-685_-decorated actin filaments. Side views of SipA_447-685_ decorated actin filaments. SipA_447-685_ subunits are colored in magenta in one groove and yellow in the other groove, and actin strands are colored in different shades of greens.

The density corresponding to SipA_447-685_ contains a compact globular domain with dimensions (30 Å x 40 Å x 40 Å) that correspond well with its previously solved crystal structure (PDB ID: 1Q5Z) (29) (Fig. 1), and a poorly resolved non-globular extension at the C-terminus that inserts deeply into F-actin (Supplementary Fig. 9). However, contrary to what was previously reported (29), we did not observe the N-terminal non-globular protrusion or arm, even though the same SipA construct was used in both studies.

### SipA binds to actin filaments with a stoichiometry of 2:4

The atomic model of the SipA-F-actin complex was built by docking the crystal structure of SipA (PDB ID: 1Q5Z) (29) and the Mg-ADP-actin filament (PDB ID: 6DJON) (34) onto the reconstructed EM map using Chimera (35) and several rounds of model building and refinement with COOT (36) and Phenix (37). Each asymmetric unit of the SipA-decorated actin filaments contains four actin subunits and two SipA subunits (Figure 2A, actin subunits are colored in light green and dark green, and SipA subunits are colored in magenta and yellow). As a result, SipA binds to actin filaments with a stoichiometry of 2:4 (Supplementary Movie 1). A globular domain that encompasses residues 514-657 and a C-terminal extension that comprises residues 658-675 of SipA could be resolved in our map (Fig. 2A). The absence of the N-terminal region (residues 447-513) in our structure suggests that this region is flexible, aligning with earlier observations that the central domain of SipA, situated in immediate proximity to the amino-terminal end of its actin-binding domain, lacks a defined structure, which was confirmed by AlphaFold modeling (31) (Supplementary Fig. 1).

**Figure 2.**
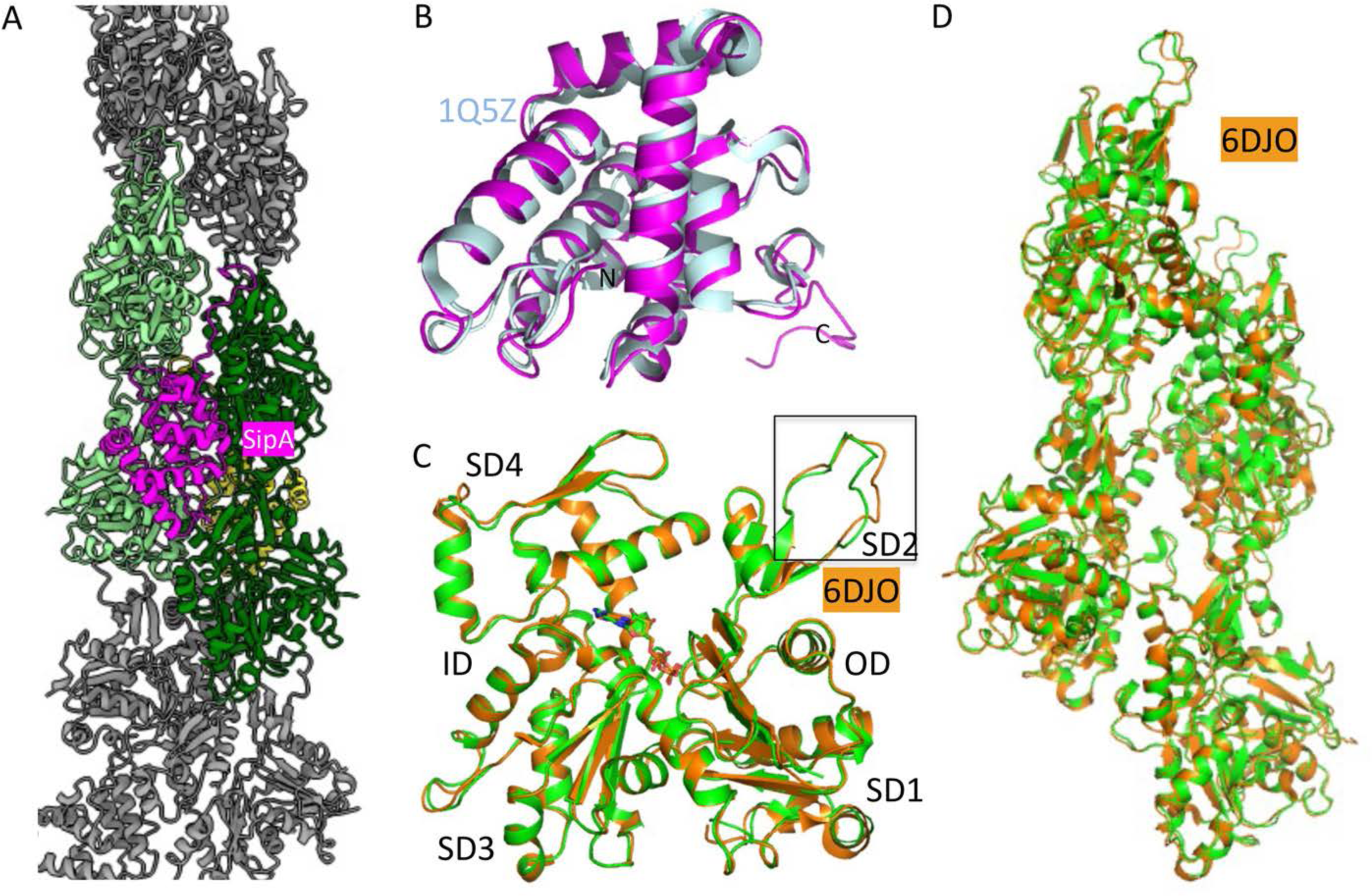
Atomic model of SipA_447-685_-decorated actin filament. (**A**) Ribbon diagram of SipA_447-685_**-**decorated actin filaments. Each asymmetric subunit contains two SipA molecules (magenta and yellow) and four actin subunits (light and dark green). (**B**) Alignment of the globular domain of SipA associated with actin filaments (magenta) to its crystal structure (PDB ID: 1Q5Z) (palecyan). (**C**) Superimposition of the structure of an actin subunit in the SipA-decorated filament to the structure of the bare actin filament (PDB ID: 6DJO). The D-loop region is highlighted with a box. The four subdomains (SD1, SD2, SD3, and SD4), inner domain (ID) and outer domain (OD) are labeled. (**D**) Alignment of actin filament in our structure to the published ADP-actin filament structure (PDB ID: 6DJO).

The alignment of the globular domain of SipA within our cryo-EM structure with its crystal structure (PDB ID: 1Q5Z) highlighted minimal conformational changes induced by actin binding, evident in a Cα root mean square deviation (RMSD) over residues 514-658 of 0.843 Å (Fig. 2B). Only minor local conformational changes in SipA were observed at its actin-binding surface. Similarly, the actin subunit in our structure and a previously solved EM structure of ADP-actin filaments (PDB ID: 6DJO) showed minimal differences (backbone RMSD: 0.610 Å) (Fig. 2C). Minor local conformational changes were observed at the actin-SipA interfaces, particularly involving the D-loop region. No significant conformational alterations were observed in the bound nucleotide, ADP, the associated cation, Mg^2+^, or the surrounding residues of the actin subunit (Fig. 2C). The actin filament in our structure aligns well with the published Mg-ADP-actin filament (PDB ID: 6DJO), suggesting that the binding of SipA does not alter the assembly of actin subunits in this filament (Fig. 2D), but only changes its helical symmetry.

In addition to the globular domain, we observed a poorly resolved carboxy-terminal extension of SipA, which inserts deeply into the actin filament (Fig. 3A and 3D and Supplementary Fig. 9). We established that, as previously shown (29), this extension is required for the ability of SipA to polymerize G-actin (Supplementary Fig. 10A and 10B) although it appears dispensable for its ability to stabilize F-actin (Supplementary Fig. 10C and 10D). Furthermore, consistent with the functional importance of this extension, its deletion also affected the ability of *S*. Typhimurium to invade cultured epithelial cells (Supplementary Fig. 10E).

**Figure 3.**
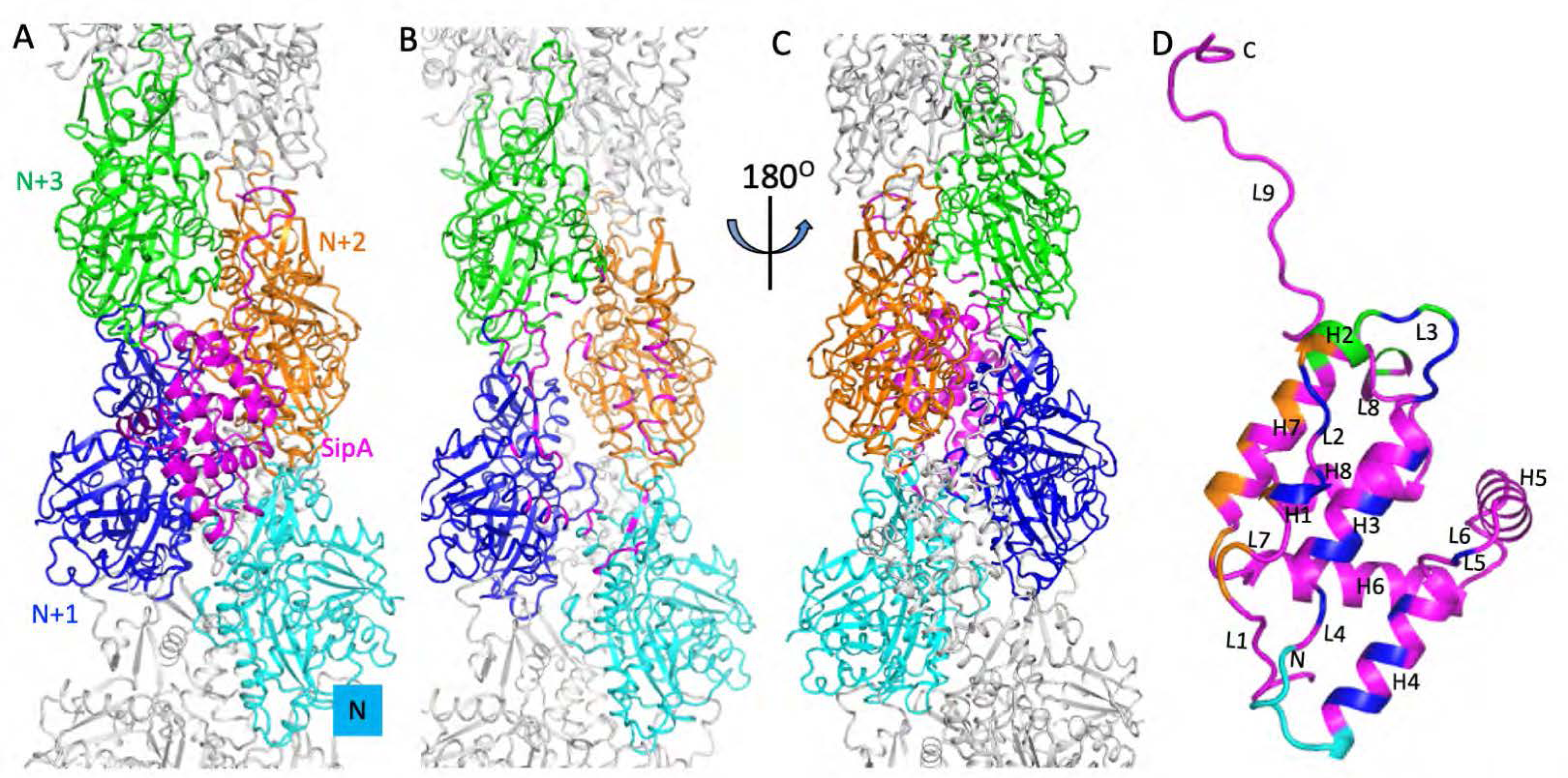
Each SipA molecule interacts with four actin subunits. (**A**) Ribbon diagram showing that one SipA_447-685_ subunit (magenta) interacts with four actin subunits (N, cyan; N+1, blue; N+2, orange; N+3, green; other subunits: gray). (**B**) SipA-binding interface on the four actin subunits. The actin residues on the actin-SipA interfaces are highlighted in magenta. SipA is hidden for clarity (**C**) Ribbon diagram showing that one SipA_447-685_ subunit (magenta) interacts with four actin subunits. The structure is rotated 180° along the filament axis relative to (**A**) and (**B**). (**D**) Actin-binding interface on SipA_447-685_. SipA_447-685_ is in the same orientation as in (**C**). Secondary structure elements are labeled, and the interacting residues are colored according to the colors of interacting actin subunits in (**C**).

### Each SipA molecule binds to four actin subunits on either side of the filament

The two strands of the actin filament create grooves running along opposite sides of the filament axis. Each SipA molecule binds to four actin subunits on either side of the two actin strands. The interface between the globular domain of SipA and the actin filament is unprecedented in that a single globular domain engages four actin subunits on both strands (N, N+1, N+2, and N+3) (Fig. 3A) through hydrophobic interactions and ionic bonds. Its interface with the actin N subunit is relatively small (∼155 Å^2^), involving residues in the SD4 domain in actin (Fig. 3B, interface residues are highlighted in magenta) and in helix 4 (H4) and loop 4 (L4) in SipA_447-685_ (Fig. 3C and 3D, interface residues are highlighted in cyan)(all the interface areas were calculated by PDBePISA). The interface between SipA and actin N+1 is large and buries ∼746 Å^2^. This interface is stabilized by more substantial interactions involving residues in the SD1 and SD2 loops of the actin subunit (Fig. 3B, interface residues are highlighted in magenta) and H1, L2, H2, L3, H3, L4, H4, and L5 in SipA_447-685_ (Fig. 3C and 3D, interface residues are highlighted in blue). The interface between SipA and actin N+2 is also substantial, burying 531.6 Å^2^ and involving actin residues mostly in the helices of SD3 and SD4 domain (Fig. 3B, interface residues are highlighted in magenta), engaging residues in L2, H1, L2, H2, L7, and H7 in SipA_447-685_ (Fig. 3C and 3D, interface residues are highlighted in orange). Finally, the interface between SipA and actin #N+3 is mainly between the loops of the SD1 and SD3 domain in actin (Fig. 3B, interface residues are highlighted in magenta), and H2, L3, H7 and L8 in SipA_447-685_ (Fig. 3C and 3D, interface residues are highlighted in green) and buries ∼214 Å^2^. Overall, the combined interface between the globular domain of SipA and F-actin is substantial thus explaining its relatively high affinity for F-actin.

### Comparison of SipA with other actin-binding molecules

Because of their affinities for polymerized actin, the fungal toxins phalloidin and jasplakinolide are frequently used to visualize actin filaments. SipA targets actin at a similar site to where these fungal toxins bind filamentous actin (Fig. 4A and 4B). However, superimposition of the structures of SipA-decorated actin filament with the toxin-actin complexes, revealed that the C-terminal extension domain of SipA would clash with the two toxin molecules, indicating the binding of SipA may block the binding of phalloidin and jasplakinolide (Fig. 4A and 4B). Pyrene labeling has been widely employed to measure the kinetics of actin polymerization and the interaction between actin and actin-binding proteins, including SipA (20, 29). Notably, pyrene binds to actin close to the SipA-binding site (38). Alignment of our structure with the structure of pyrene-bound actin filament (PDB ID:7K21) shows that the binding of pyrene alters the conformation of the D-loop region of the actin subunit in a manner that would clash with SipA. More specifically, V45 on the D-loop region of actin clashes with H539 of SipA (Fig. 4C). These findings therefore suggest that actin polymerization assays employing pyrene actin may not be suitable for studying the effect of SipA on actin dynamics.

**Figure 4.**
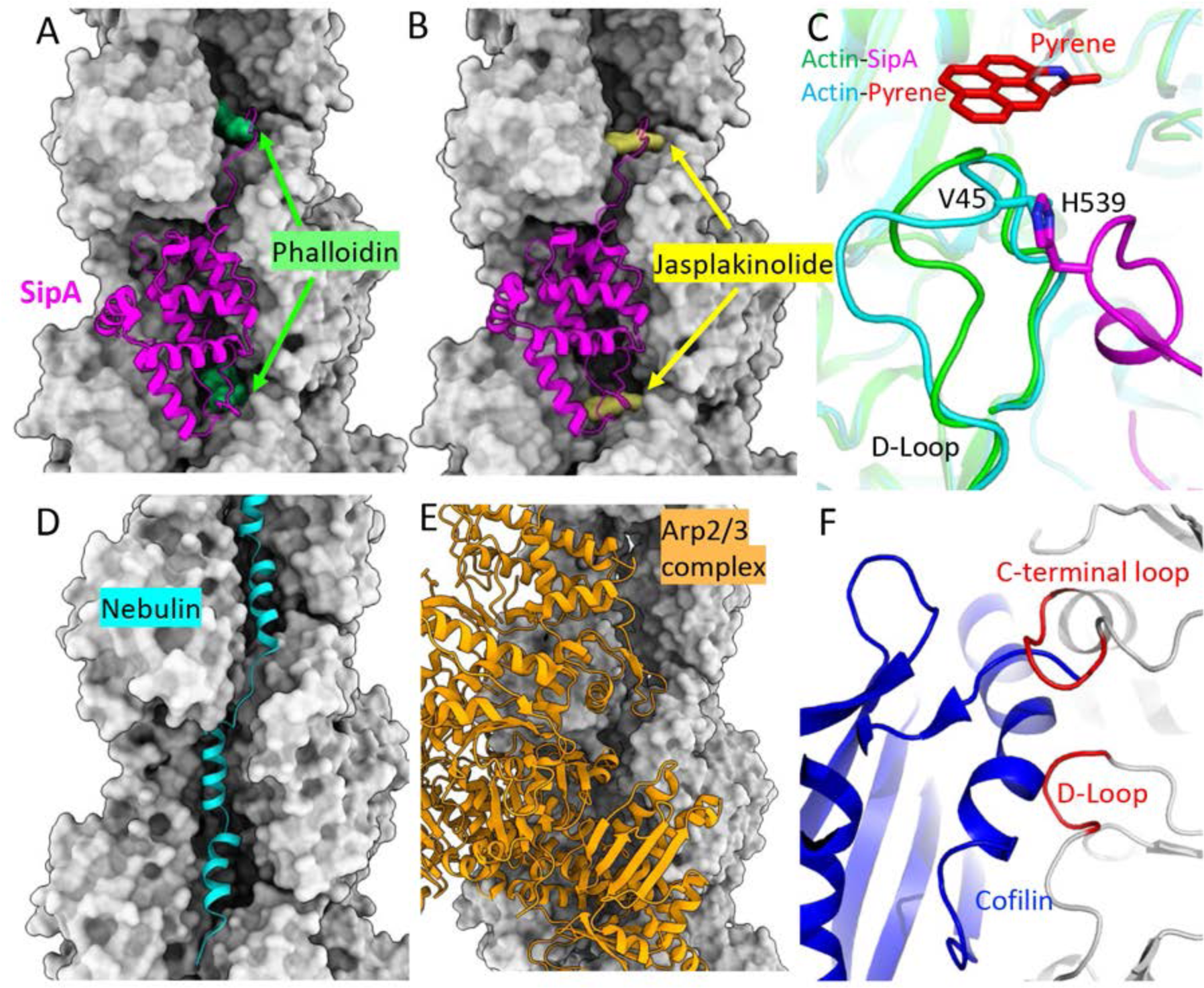
Comparison of the actin-binding interface of SipA with other actin binding molecules. (**A-C**) Superimposition of the structures of SipA and phalloidin (PDB ID: 7BTI) (**A**), jasplakinolide (PDB ID: 6T23) (**B**), or pyrene (PDB ID: 7K20) (**C**) bound to actin. (**D**) Structure of Nebulin (PDB ID: 7QIM) bound to actin filaments showing that nebulin and SipA bind to the same groove of the actin filament. (**E**) Structure of the Arp2/3 complex bound to actin filaments (PDB ID: 8E9B) depicting that SipA shares the same binding site. (**F**) Cofilin clashes with the actin D-loop region and the C-terminal loop (S348-F352) in the SipA-decorated actin filament. The inner domain (subdomains 3 and 4) of an actin subunit is used for structure superimposition. SipA, magenta; actin filament, light gray; phalloidin, green; jasplakinolide, yellow; nebulin, cyan; Arp2/3 complex, orange; cofilin, blue.

The actin-binding site of SipA also overlaps with the binding site of other well-characterized actin-regulatory proteins. For example, SipA binds the same groove in F-actin that serves as a binding site for the muscle protein nebulin (Fig. 4D) (39, 40). Based on low-resolution structures of SipA and Nebulin bound to actin, it was previously proposed that SipA mimics nebulin in its mechanism of actin-binding (30). Our structure is consistent with this notion since several actin residues engaged by SipA (e.g. K68, E117, E224, E276, and R372) are also engaged by nebulin (41).

The Arp2/3 complex forms branches by nucleating the daughter filament from the side of the mother actin filaments (42). Superimposition of our structure and the previous structure of the actin branch junction (PDB ID: 8E9B) (43) shows that SipA and Arp3 bind at the same site on actin filaments (Fig. 4E). Thus, binding of SipA likely inhibits the binding of Arp2/3 complex thus potentially preventing the branching of actin filaments.

It was previously reported that SipA interferes with the F-actin severing activity of ADF/cofilin (25). Although SipA and cofilin (44) bind actin at different sites, the SipA-bound filaments have different helical symmetries therefore potentially interfering with cofilin binding. In fact, superimposing the SipA- and cofilin-bound (5YU8) filamentous actin structures using the inner domain (subdomains 3 and 4) of one actin subunit shows that Cofilin would clash with the D-loop region and the C-terminal loop (S348-F352) of actin in its SipA-bound configuration (Fig. 4F). These observations provide a plausible mechanistic explanation for the observation that actin filaments bound to SipA are resistant to the severing activity of ADF/cofilin (25).

### Bacterial actin homologs lack structural features that are essential for SipA binding

Many bacteria, including *Salmonella*, encode highly conserved cytoskeletal proteins such as MreB and ParM, that are structural homologs of actin (45, 46). However, despite the close structural similarities between mammalian actin and the bacterial counterparts, SipA should have evolved to be unable to target these bacterial actin homologs as such binding would likely be detrimental to bacterial physiology and could potentially trap SipA within the bacterial cytoplasm. Consistent with this notion, comparison of the structures of MreB and ParM with actin shows that several actin structural features that are crucial for the formation of its interface with SipA are partially or totally absent from MreB and ParM (Fig. 5). These results indicate that, as previously suggested (30), SipA has evolved to avoid binding to the bacterial actin homologs.

**Figure 5.**
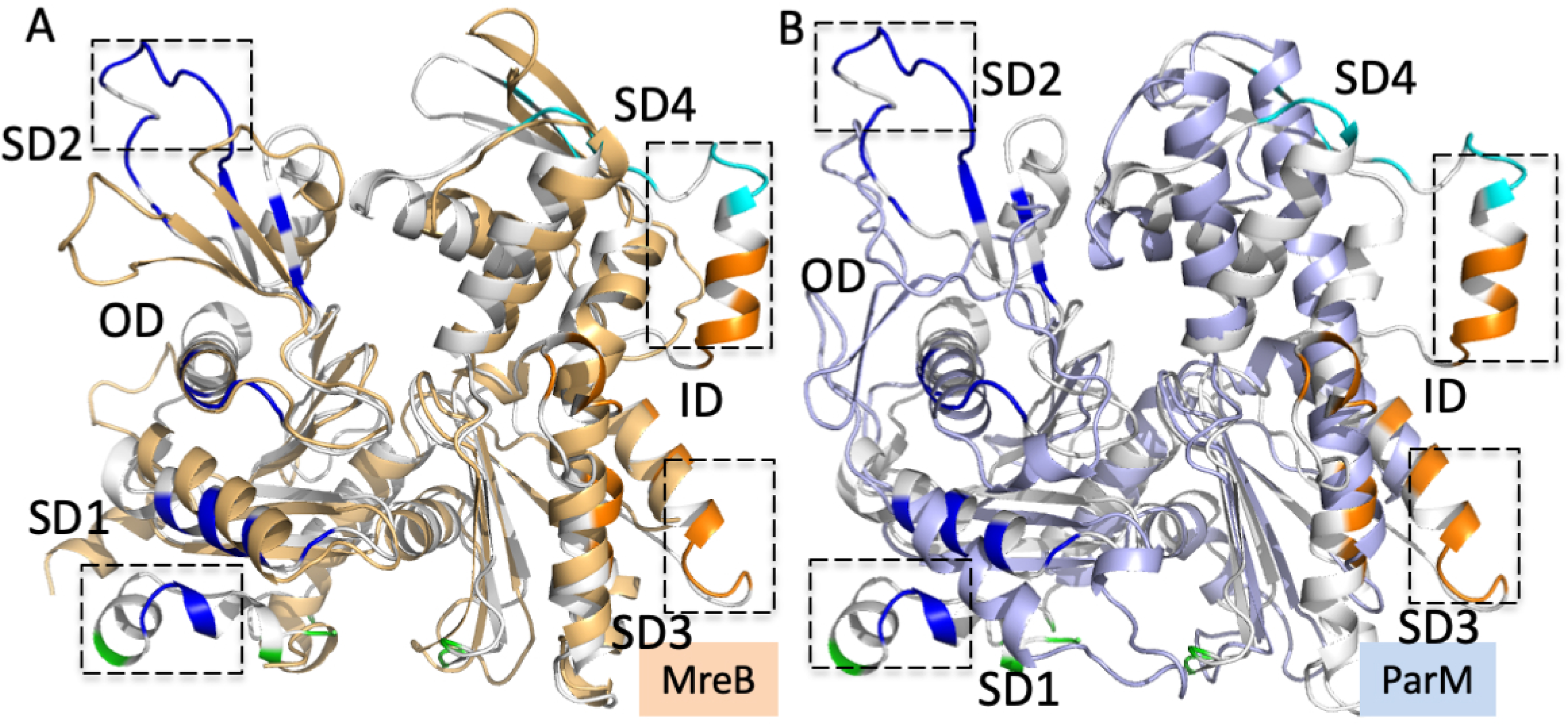
SipA binding sites are absent from the bacterial actin homologs MreB and ParM. (**A**-**B**) Alignment of MreB (tan) (**A**) or ParM (light purple) (**B**) to an actin subunit bound to SipA (white). Residues at different SipA-actin interfaces are highlighted following the same color scheme as in Figure 3A. The loops and helices of actin not present in ParM or MreB but involved in the SipA-actin interactions are highlighted. The four subdomains (SD1, SD2, SD3, and SD4), inner domain (ID) and outer domain (OD) of actin subunit are labeled.

### Structure-function analyses of SipA indicate that its actin-modulating and pro-inflammatory activities are independent of one another

SipA has been associated with two different phenotypes, bacterial invasion of intestinal epithelial cells through its actin-modulating activity (20), and the stimulation of intestinal inflammation through poorly understood mechanisms that involve its amino-terminal half (26, 27, 47). Whether the ability of SipA to stimulate intestinal inflammation requires its actin-modulating activity is unclear since the lack of structural information has prevented the generation of a mutant specifically defective in this function while preserving the proper folding and integrity of the entire protein. This issue is pertinent since it has been shown that actin polymerization, when triggered by microbial pathogens, can activate innate immune signaling (48). To address this issue, we introduced mutations in SipA residues that are implicated in the formation of its interfaces with actin and examined their effect on F-actin-binding and G-actin polymerization. Reflecting the redundant nature of the SipA-actin interface, none of the individual mutations introduced significantly affected the ability of SipA to bind actin (Fig. 6A). However, introduction of changes in two or more SipA residues involved in forming its interface with actin significantly reduced its binding affinity. Notably, simultaneous mutations in S537 and K636, which interact with the D-loop and C-terminus of actin (Supplementary Fig. 11), resulted in a drastic reduction in its binding affinity (Fig. 6A).

**Figure 6.**
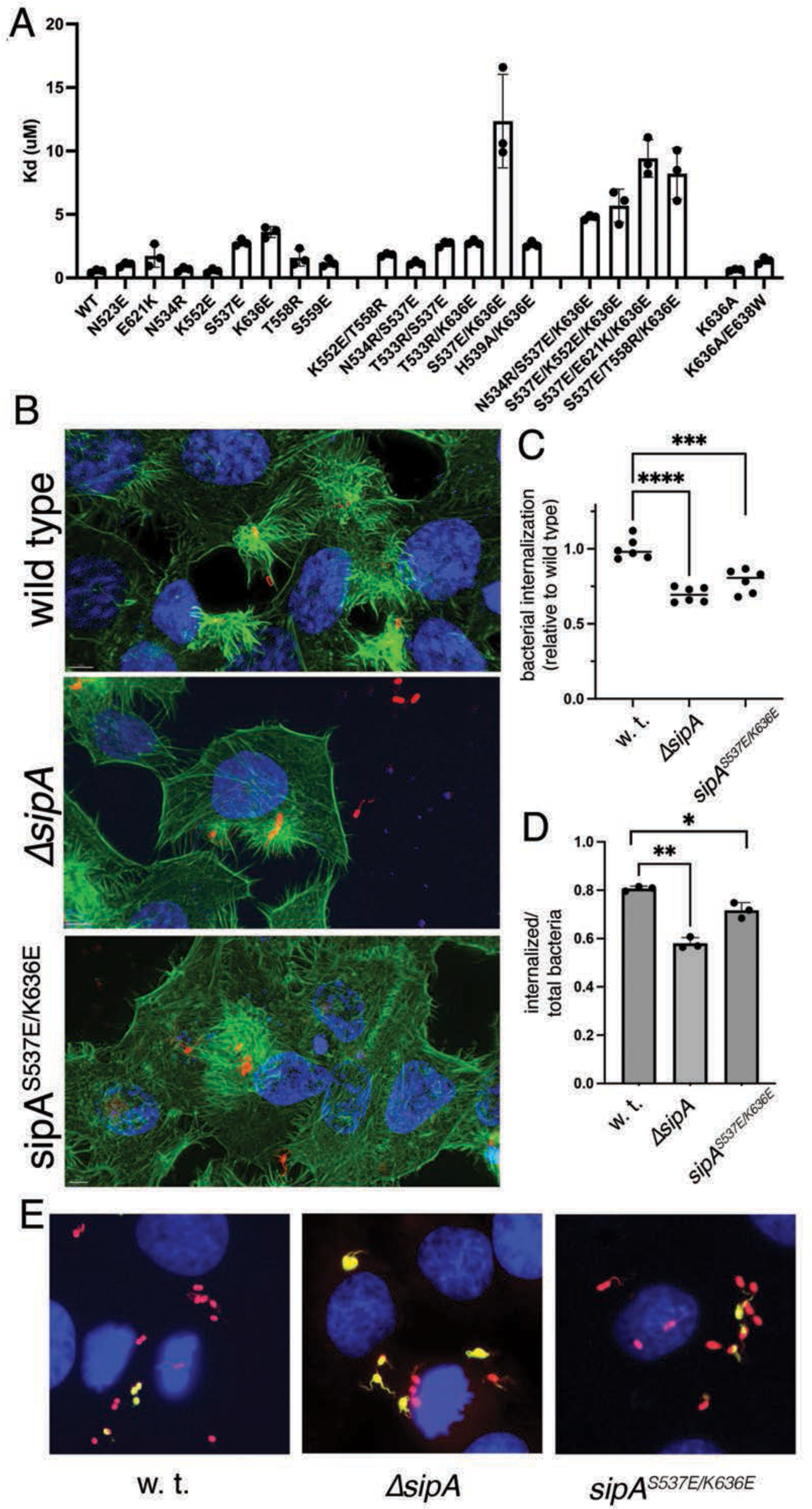
Structure-function analysis of the SipA-actin interaction. (**A**) Determination of the binding affinity of different SipA constructs to actin filaments with a co-sedimentation assay. The SipA mutations are on their binding interfaces. The bars are the average dissociation constants calculated from three independent experiments, and the errors are the standard deviations. (**B**) Actin cytoskeletal rearrangements observed after infection with wild-type *S*. Typhimurium (left), a *ΔsipA* isogenic mutant (middle), and a strain expressing the SipA^S537E/K636E^ mutant (right), which showed decreased binding affinity to actin. Actin filaments were stained with Atto 488-phalloidin (green), bacteria were stained with an antibody directed to the *S*. Typhimurium LPS (red), and mammalian cell nuclear DNA was stained by 4’,6-diamidino-2-phenylindole (DAPI) (blue). (**C**-**E**) Cultured epithelial cell invasion of wild type *S*. Typhimurium, its isogenic *ΔsipA* mutant or a mutant expressing SipA^S537E/K636E^, which is defective for actin binding. The invasion ability of different strains was measured by the gentamicin protection (**C**) or differential (inside-out) staining (**D** and **E**) assays. Values in (**C**) are the ratio of the inoculum that survived antibiotic treatment due to internalization, normalized to wild type (w. t.), which was set to 1. The mean was calculated from six experiments (***: P<0.001; ****: P<0.0001, two-tailed Student *t*-test). w. t.: wild type. (**D**) Quantification of internalized bacteria by the differential staining assay. Values represent mean ± SD of the ratio of internalized vs total bacteria derived from quantifying a minimum of 1,000 cells per condition. (*: P<0.05; **: P<0.01, two tailed Student *t* test). (**E**) Representative fluorescence microscopy images showing stained nonpermeabilized (green) or all (nonpermeabilized and permeabilized, red) *S*. Typhimurium after infection. The mammalian cell nuclear DNA was stained by 4’,6-diamidino-2-phenylindole (DAPI) (blue).

To examine the contribution of the actin-binding activity of SipA to the interaction of *S.* Typhimurium with host cells, we constructed a mutant strain expressing the *sipA^S537E/K636E^* allele from its native chromosomal context. We then tested the resulting strain’s ability to stimulate actin-cytoskeletal rearrangements and invade cultured epithelial cells. We found that, in comparison to the wild type, the actin cytoskeletal changes stimulated by the *S.* Typhimurium *sipA^S537E/K636E^* mutant strain were more diffuse, with less defined ruffles (Fig. 6B). Consistent with the more limited actin rearrangements, the invasion ability of the mutant strain was significantly impaired when measured with two independent assays (Fig. 6C-6E). Notably, the phenotype of the *S.* Typhimurium *sipA^S537E/K636E^* mutant strain was similar to the phenotype of the *S*. Typhimurium *ΔsipA* mutant, further demonstrating that the introduced mutations largely eliminated its ability to bind actin (Fig. 6C-6E).

To investigate the role of SipA’s actin-modulating activity in promoting inflammation, we compare the intestinal inflammatory response of mice infected with different *S*. Typhimurium strains. Specifically, we compared mice infected with a strain lacking *sipA*, a strain expressing the actin-binding defective mutant SipA^S537E/K636E^, and a strain expressing wild-type SipA. To enhance the detection of SipA-dependent inflammation (27), we introduced a *ΔspiA* mutation in all strains, rendering them deficient for the SPI-2 T3SS. We found that mice infected with *S*. Typhimurium expressing either wild type SipA or the actin-binding defective mutant displayed smaller cecum sizes and lighter weights indicative of reduced cecal content due to inflammation (Fig. 7A and 7B). In addition, these mice exhibited substantial intestinal inflammation, as observed by histopathology (Fig. 7C). In contrast, mice infected with the *sipA*-deficient *S*. Typhimurium strain exhibited cecum sizes similar to uninfected controls (Fig. 7A and 7B) and much reduced intestinal inflammation (Fig. 7C). These results indicate that the actin-binding activity of SipA is not necessary for inducing intestinal inflammation. Thus, the actin-modulating and pro-inflammatory activities of SipA are independent of each other.

**Figure 7.**
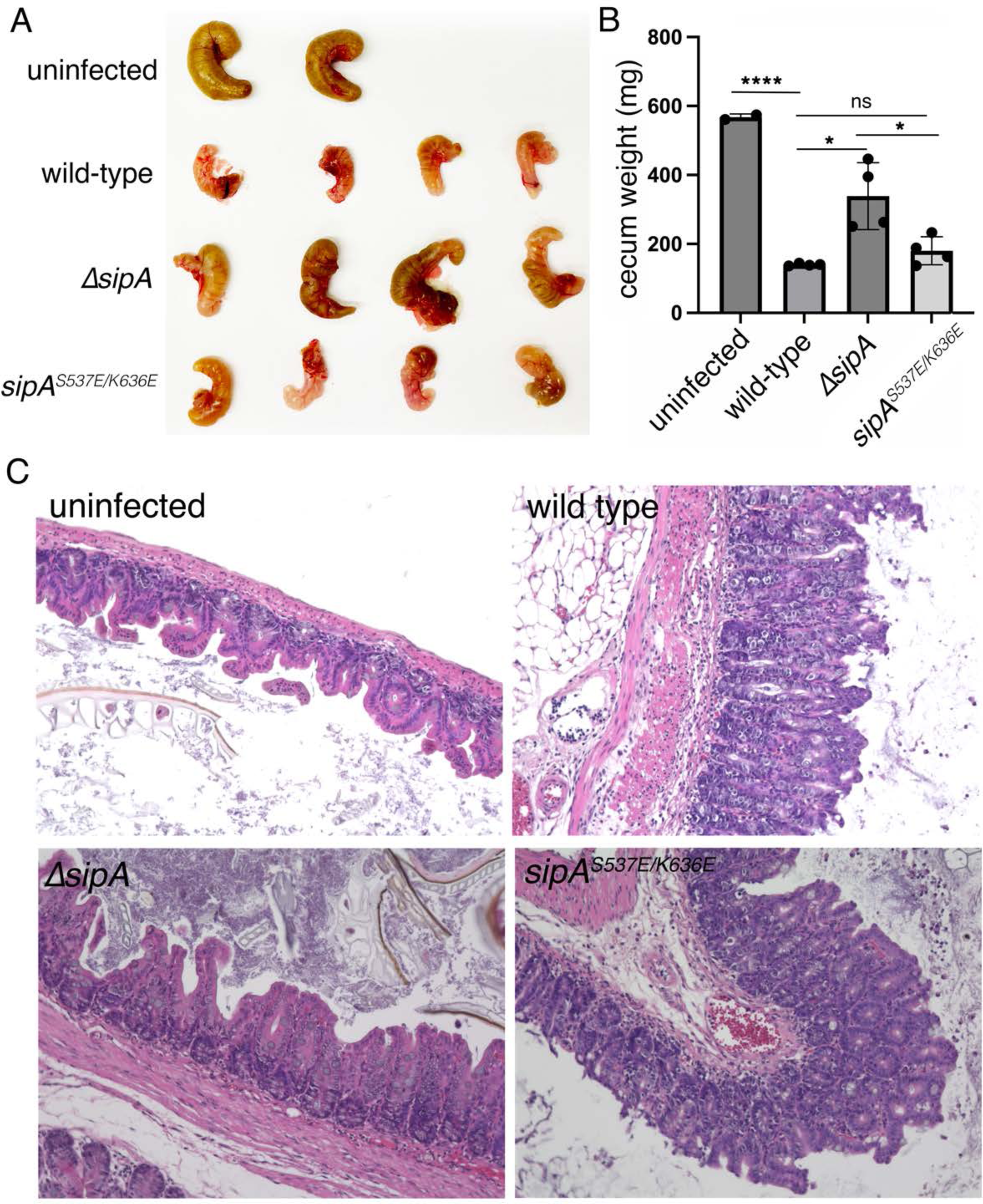
The pro-inflammatory activity of SipA is independent of actin-binding. Mice were infected with *S*. Typhimurium strains lacking *sipA*, expressing the actin-binding defective mutant SipA^S537E/K636E^, or expressing wild-type SipA. To enhance the detection of SipA-dependent inflammation (27), a *ΔspiA* mutation was introduced in all strains, rendering them deficient for the SPI-2 T3SS. Gross pathology (**A**), weight (**B**), and histopathology (**C**) of ceca obtained from mice infected with the different strains or uninfected controls, as indicated, are shown. Values in (**B**) are the weight of each cecum in mg and represent the mean ± SD (*: P<0.05; ****: P<0.0001; ns: not signifcant; two tailed Student *t* test)

## Discussion

Bacterial effector proteins that can modulate cellular processes are often evolutionary designed to mimic the activities of eukaryotic proteins that carry out a similar function (49, 50). This mimicry often involves convergent evolution since many of these bacterial mimics do not exhibit either primary sequence or structural similarity to their eukaryotic counterparts (49). The actin-binding effector protein SipA has been postulated to mimic the actin-binding protein nebulin (30). In addition, it has been proposed to exert its actin-stabilizing function by working as a “molecular staple”, using two flexible arms to bind actin subunits on opposing actin strands (29). However, the lack of a high-resolution structure of SipA bound to actin filaments has prevented the rigorous testing of these hypotheses. To address these issues, we solved the cryo-EM structure of the actin-binding carboxy-terminal domain of SipA bound to actin filaments. We found that each SipA molecule simultaneously engages four actin subunits through interactions involving only its globular carboxy-terminal domain. The SipA binding sites on actin overlap those of nebulin, a very large muscle skeletal protein, which through the tandem arrangement of multiple actin-binding domain repeats can engage up to 200 actin subunits (39). Our structure showed that SipA and each nebulin repeat bind the same groove in the actin filaments and share some actin residues necessary for their interaction. This observation is therefore consistent with the notion that SipA and the nebulin repeat exhibit a similar actin-binding mode.

Our structure also showed that the SipA actin-binding site overlaps with the binding site of a well-characterized actin nucleator, the Arp2/3 complex. This observation has potential functional relevance since it suggests that SipA is likely to interfere with the binding of the Arp2/3 complex, thus potentially preventing the branching of SipA-stabilized actin filaments. However, more experiments will be required to test this hypothesis and to examine its potential functional significance.

It has been previously reported that SipA prevents the activity of the actin-severing proteins ADF/cofilin (25), resulting in further stabilization of SipA-bound filaments, enhanced membrane ruffles and bacterial internalization. Our structure provides a plausible mechanistic explanation for such activity. The actin-binding sites of SipA and ADF/Cofilin do not overlap (44). However, SipA-bound filaments exhibit a different helical symmetry from those bound by cofilin, which likely interferes with its binding. In fact, comparison of the structures of SipA- and cofilin-bound filaments predicts clashes between the D-loop region and the C-terminal loop (S348-F352) of actin in its SipA-bound configuration and cofilin potentially explaining the inhibitory activity.

Our structure of SipA bound to actin did not reveal the presence of non-globular N-terminal arms engaging actin subunits on the opposite strand of the actin filaments, as observed in a previous low-resolution cryo-EM structure (29). Thus, we could not confirm the “molecular staple” model. However, our structure did show an unstructured extension at the C-terminal end, which inserts deep into the actin filament. Furthermore, as previously shown(29), we found that this extension is required for the ability of SipA to increase the efficiency of bacterial invasion of cultured epithelial cells. This extension most likely contributes to bacterial invasion by facilitating the polymerization of G-actin, rather than by stabilizing F-actin, as removing this extension affected the former but not the latter.

In addition to its ability to modulate actin dynamics to mediate bacterial invasion, SipA has also been implicated in the stimulation of intestinal inflammation (26, 27), which is critical for *S*. Typhimurium virulence (51). Although the mechanisms by which SipA modulates the inflammatory response are poorly understood, this activity has been largely correlated with its amino-terminal domain (26, 27, 47). However, as the modulation of actin dynamics by bacterial pathogens has also been linked to the stimulation of innate immune responses involving inflammation (48), it has been unclear whether the carboxy-terminal, actin-binding domain of SipA contributes to this phenotype. Indeed, the lack of structural information for the SipA-actin interface has hampered previous efforts to obtain a mutant that would specifically lack this activity without altering the overall structure of this effector protein (28). This issue is complicated by the redundant nature of the SipA-actin interface. In fact, a previously reported mutant utilized in a study to attempt to address this issue (28) in our hands retains the ability to modulate actin dynamics (Fig. 6). Using structurally guided mutagenesis, we were able to identify a mutant that is specifically defective in actin binding without altering the overall conformation of SipA. As predicted by its significantly decreased ability to bind actin, an *S*. Typhimurium strain expressing this mutant stimulated more diffuse actin cytoskeleton rearrangements, less prominent ruffles, and was defective in bacterial invasion of cultured epithelial cells. In fact, these phenotypes were similar to those of a *S*. Typhimurium strain lacking *sipA*. To address the potential contribution of the actin-binding activity to the stimulation of intestinal inflammation, we tested the *S*. Typhimurium strain expressing the actin-binding defective SipA mutant for its ability to stimulate intestinal inflammation in a mouse model of infection. Surprisingly, we found that the *S.* Typhimurium strain expressing the SipA mutant was able to stimulate increased inflammation in comparison to the wild type. These results indicate that the actin-binding activity of SipA does not contribute to its ability to induce inflammation.

Taken together, our structure function studies have revealed a unique mechanism by which a bacterial effector protein modulates actin dynamics. Indeed, the ability of a protein to bind four actin subunits with a single globular domain has not been previously reported for other actin-binding proteins. Furthermore, our studies have clarified the mechanism of action of this effector protein and provide unique insight into the *Salmonella* spp. virulence mechanisms.

## Materials and Methods

### Bacterial strains and growth conditions

The wild-type *Salmonella enterica* serovar Typhimurium SL1344 strain and its *ΔsipA* or ΔspiA (SPI-2 T3SS deficient) isogenic mutants have been previously described (20, 52). The *S*. Typhimurium SL1344 derivative encoding SipA^S537E/K636E^ was constructed by allelic exchange as previously described (53). All bacterial strains were cultured in L-Broth containing 0.3 M NaCl to an OD600 of 0.9, conditions known to stimulate the expression of the SPI-1 T3SS (54).

### Plasmid construction, protein expression, and purification

DNA manipulations were carried out using the standard Gibson assembly protocol (55). *S.* Typhimurium SipA_447-685_ was expressed in a modified pQE vector with GST and a 6xHis tag at its amino and carboxy-terminal ends, respectively. SipA_447-685,_ SipA_498-659_, and SipA_513-659,_ as well as SipA_447-685_ carrying specific point mutations, were expressed in *Escherichia coli* BL21 (DE3) and purified using affinity chromatography. Briefly, bacterial cells were cultured in LB medium at 37°C to an absorbance at OD_600_ of 0.8 to 1.0, induced with 0.2 mM isopropyl β-d-1-thiogalactopyranoside (IPTG) and grown for an additional 16 to 20 h at 20°C. The bacteria were harvested by centrifugation and lysed using a One-Shot cell disrupter following the manufacturer’s instructions (Constant Systems Ltd., Daventry, United Kingdom). Cell lysates were cleared by centrifugation, and the supernatant was loaded onto a Glutathione Sepharose 4B gravity-driven column. After elution, the GST tag was removed by digestion with HRV 3C protease. The digested protein mixture was further purified by passing through a second Glutathione Sepharose 4B and a Ni^2+^–nitrilotriacetic acid column.

Actin was purified as previously reported (56). Briefly, actin was purified from chicken muscle acetone powder made from flash-frozen chicken (purchased from a local Trader Joe’s store). After one cycle of polymerization and depolymerization, G-actin was isolated by gel filtration through Sephacryl S-300 column in Ca-G-buffer (2 mM Tris⋅HCl, pH 8.0; 0.2 mM ATP; 0.1 mM CaCl_2_; 1 mM NaN3; 0.5 mM DTT).

### Cell culture, bacterial infections, and visualization of the actin cytoskeleton

All cell lines were routinely tested for the presence of mycoplasma by a standard PCR method, checked for their morphological features, growth characteristics and functionalities, but were not authenticated by short tandem repeat profiling. The ability of *S*. Typhimurium strains to invade cultured human intestinal epithelial Henle-407 cells (obtained from the Roy Curtiss III collection in 1987) using the gentamicin protection assay was carried out as described previously (57). Briefly, ∼24 h after seeding onto tissue culture plates, cells were infected for 20 min with the indicated *S*. Typhimurium strains with an m.o.i. of 20. Infected cells were washed and incubated for 2 h in DMEM containing gentamicin (100 µg ml^−1^) to kill extracellular bacteria. Alternatively, bacterial invasion was also measured using a differential staining protocol (inside-out staining) as described before (58). Briefly, cells were infected as indicated above and 20 min after infection, the cells were fixed in formalin and stained with rabbit anti-*Salmonella* lipopolysaccharide (LPS) antibody (generated at Poconos farms), followed by fluorescently labeled (Alexa Fluor 488) secondary anti-rabbit antibody. Cells were subsequently permeabilized with Triton X 100 (0.3 %) and stained with the anti-Salmonella LPS antibody followed by a fluorescently labeled (Alexa Fluor 594) anti-rabbit secondary antibody. In this protocol, extracellular bacteria are stained by both secondary antibodies, while intracellular bacteria are only stained by the Alexa Fluor 594-labeled secondary antibody. Internalized bacteria were quantified by capturing a minimum of 30 fields per condition (containing a minimum of 1000 cells each) on a Nikon Eclipse inverted microscope equipped with an Andor’s Zyla 5.5 sCMOS camera.

To visualize actin cytoskeleton rearrangements due to bacterial invasion, cells were infected, fixed, and permeabilized as indicated above and then stained with Atto-488-labeled phalloidin (to visualize the actin cytoskeleton), an anti-*S.* Typhimurium LPS antibody (SIFIN, catalog no.: TS 1624), and Alexa Fluor 594 fluorescently labeled secondary antibody to visualize bacteria. Cells were visualized under an Andor Dragonfly spinning disk fluorescence microscope (Oxford Scientific).

### Mouse infections and measurement of intestinal inflammation

Groups of age- and sex-matched C57BL/6 mice were orally administered streptomycin (100 µl of 200 mg/ml solution) and 24 hrs after antibiotic treatment were orally infected by stomach gavage with 10^7^ cfu in 100 μl of PBS with the indicated strains. Forty-eight hours after infection, mice were euthanized and the cecum was collected. Ceca were weighted as a proxy measurement of inflammation and then fixed in formalin and embedded in paraffin for sectioning and H&E staining. Embedding, sectioning, and staining were performed by the Yale Pathology Tissue Services. The sample sizes were empirically determined to optimize the numbers based on our previous experience with equivalent experiments. The mice were randomly assigned to the experimental groups, but experimenters were not blind to the assignment.

### EM sample preparation, data acquisition, and image processing

After cation exchange from Ca^2+^ to Mg^2+^ using 0.1 volumes of 10×ME buffer (0.5 mM MgCl_2_, 2 mM EGTA, pH7.5), Mg-ATP-actin monomers were first polymerized in KMEI buffer (50 mM KCl, 1 mM MgCl_2_, 1 mM EGTA and 10 mM Imidazole pH 7.0) for 20 min, followed by the addition of SipA with a molar ratio of 1.7:1. The CapZ protein was added into the polymerized actin filaments with a molar ratio of 1:30 to avoid the formation of long actin filaments. For negative staining, 3.5 µl of actin filaments (final concentration: 3 µM) or SipA (final concentration:5 µM)-decorated actin filaments (final concentration: 3µM) were applied onto glow-discharged carbon-coated grids and negatively stained with 0.8% (w/v) uranyl formate. Images were collected using a Talos L120C electron microscope (Thermo Fisher) operating at 120 kV.

For cryo-EM analysis, 3 µl of the sample was deposited onto glow-discharged Quantifoil R2/1 300 Mesh Gold Holey Carbon Grids (SPI supplies) and plunged into liquid ethane using a Vitrobot Mark IV (Thermo Fisher) at 10°C and 100% humidity. The freezing conditions were screened on a Glacios cryo-TEM (Thermo Fisher) operating at 200 kV. Cryo-EM data collection was performed using a Titan Krios electron microscope (Thermo Fisher) equipped with a K3 direct electron detector (Gatan) operating at 300 kV. Data were collected at a calibrated magnification of 81,000x (1.068 Å per pixel) in super-resolution mode, with defocus values between −2.5 µm and −1.2 µm. Images were recorded using the beam image shift implemented in SerialEM (59). Forty-four frames were taken at 0.05s per frame, with a dose rate of 29 e^−^ per pixel per second. In total, 4740 movies were recorded with an exposure time of 2.2 s. Motion correction and contrast transfer function (CTF) estimation were done using Relion 4.0 (60). Filaments were manually picked by using sxhelixboxer.py in SPARX (61). Layer-line images were calculated with SPARX (project and periodogram commands). In total, 278,740 filament segments were boxed out from 2,370 images and subjected to further 2D and 3D calculations in Relion 4.0. The cryo-EM map of ADP-actin (EMDB accession number: 7938) was used as the initial map and helical symmetry parameters (rise: 109.9 Å and twist: 53.3 ^O^) were imposed on the reconstruction. 3D classification resulted in classes with different SipA occupancies. A class with most particles were selected and subjected to additional round of 3D classification. Three classes were combined (69,483 particles) for further 3D refinement. DeepEMhancer (33) was employed to improve map connectivity and interpretability. The final map has a global nominal resolution of 3.6 Å (based on the FSC using the 0.143 criterion calculated in Relion 4.0). The local resolution was calculated with ResMap in Relion 4.0. All the structural calculations were carried out on the Yale High-Performance Computing servers. The statistics for data collection and map reconstruction are summarized in Table S1. The data processing workflow is described in Supplementary Fig. 12.

### Model Building and refinement

The cryo-EM structure of Mg-ADP-actin filaments (PDB ID: 6DJO) (38) and the crystal structure of SipA (PDB ID: 1Q5Z) (29) were used as initial templates to dock into the EM maps using Chimera (35). The modeled structure was manually adjusted in Coot (62) and refined with Phenix (63). The SipA-actin interfaces were calculated with PDBePISA (64). Figures were generated using PyMOL and Chimera. The statistics for model refinement are listed in Table S1.

### Actin co-sedimentation assay

To test the binding affinity of the different SipA mutants, actin monomers were first polymerized into filaments in KMEI buffer (50 mM KCl, 1 mM MgCl2, 1 mM EGTA and 10 mM Imidazole pH 7.0) for 20 min. Then different concentrations (0, 1, 2, 3, 4, and 10 µM) of polymerized actin filaments were incubated with 4 µM of wild-type SipA or SipA mutants for 10 min at room temperature. Followed by ultracentrifugation at 120,000 x g in a TLA100 rotor (Beckman Instruments, Fullerton, CA) at 4°C for 30 min, an equal amount of the supernatant from each reaction was loaded onto SDS–PAGE Gel and stained with Coomassie blue.

To test the actin nucleation activity of SipA_447-685_, SipA_498-659_, and SipA_513-659_, the SipA proteins were first dialyzed against a Tris buffer without salt (25 mM Tris, pH 7.5) three times before adding to G-actin. After 4 hs of incubation, the samples were ultracentrifuged at 120,000 x *g* in a TLA100 rotor (Beckman Instruments, Fullerton, CA) at 4°C for 30 min, then an equal amount of the supernatant and pellet from every reaction were loaded onto the SDS–PAGE gel and stained with Coomassie blue.

### Data Availability

The EM maps and atomic coordinates have been deposited in Electron Microscopy Data Bank (EMDB) under accession number EMD-43188 and Protein Data Bank (PDB) under accession code 8VFM.

### Ethics statement

All animal experiments were conducted according to protocols approved by Yale University’s Institutional Animal Care and Use Committee (IACUC) under protocol number 2022–07858. The IACUC is governed by applicable Federal and State regulations, including those of the Animal Welfare Act, Public Health Service and the United States Department of Agriculture and is guided by the US Government Principles for the Utilization and Care of Vertebrate Animals Used in Testing, Research and Training.

## Acknowledgements

We thank Dr. Jianfeng Lin, Dr. Shenping Wu, Dr. Marc Llaguno, and Dr. Xinran Liu for managing the microscope resources, the Yale High Performance Computing support team for guidance and use of the computing infrastructure. We also thank Florian Schueder for assistance in the imaging of the actin cytoskeleton rearrangements induced by the different *S*. Typhimurium strains. This work was supported by NIH Grant R01AI055472 to J. E. G.

## Supplementary Figures for

**Supplementary Fig. 1.**
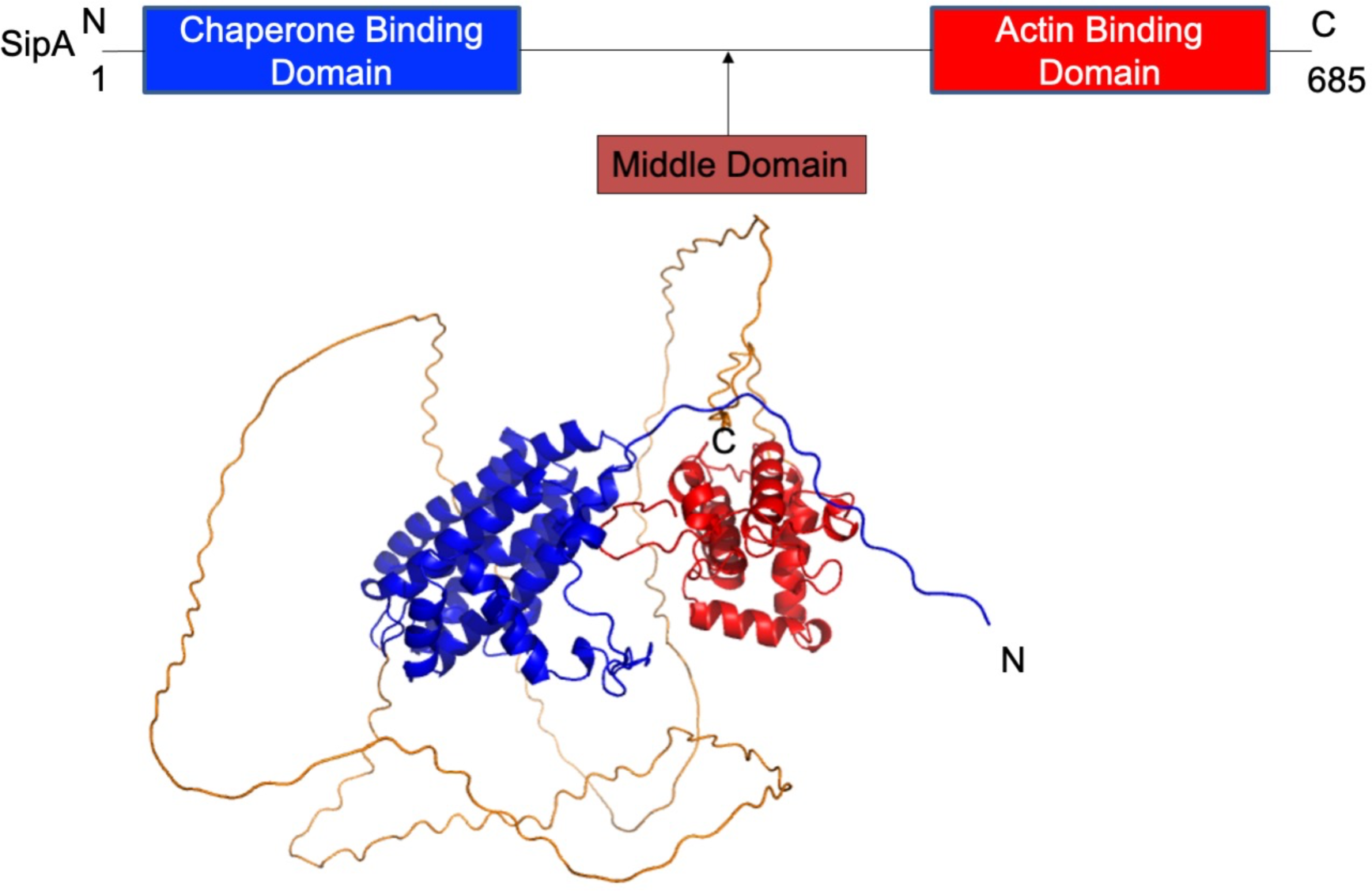
Diagram of SipA and AlphaFold prediction of its structure.

**Supplementary Fig. 2.**
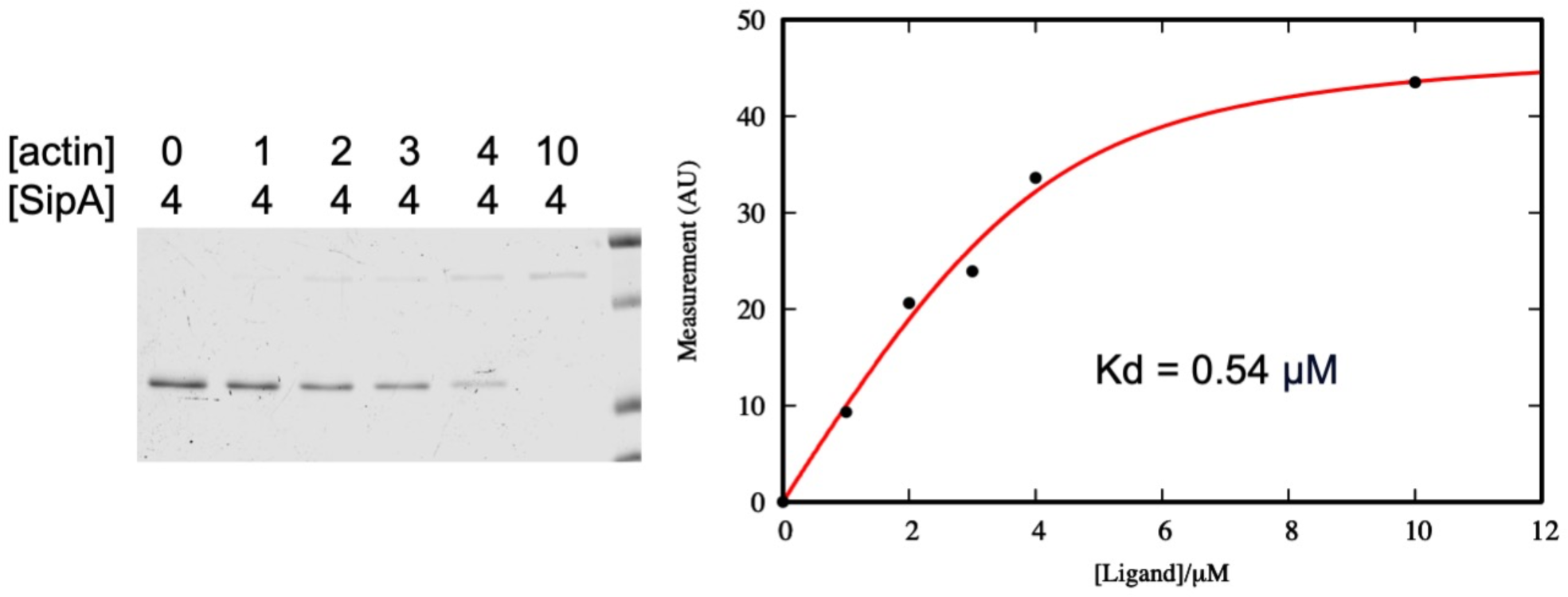
Actin-binding affinity of SipA_447-685._ as determined by a co-sedimentation assay.

**Supplementary Fig. 3.**
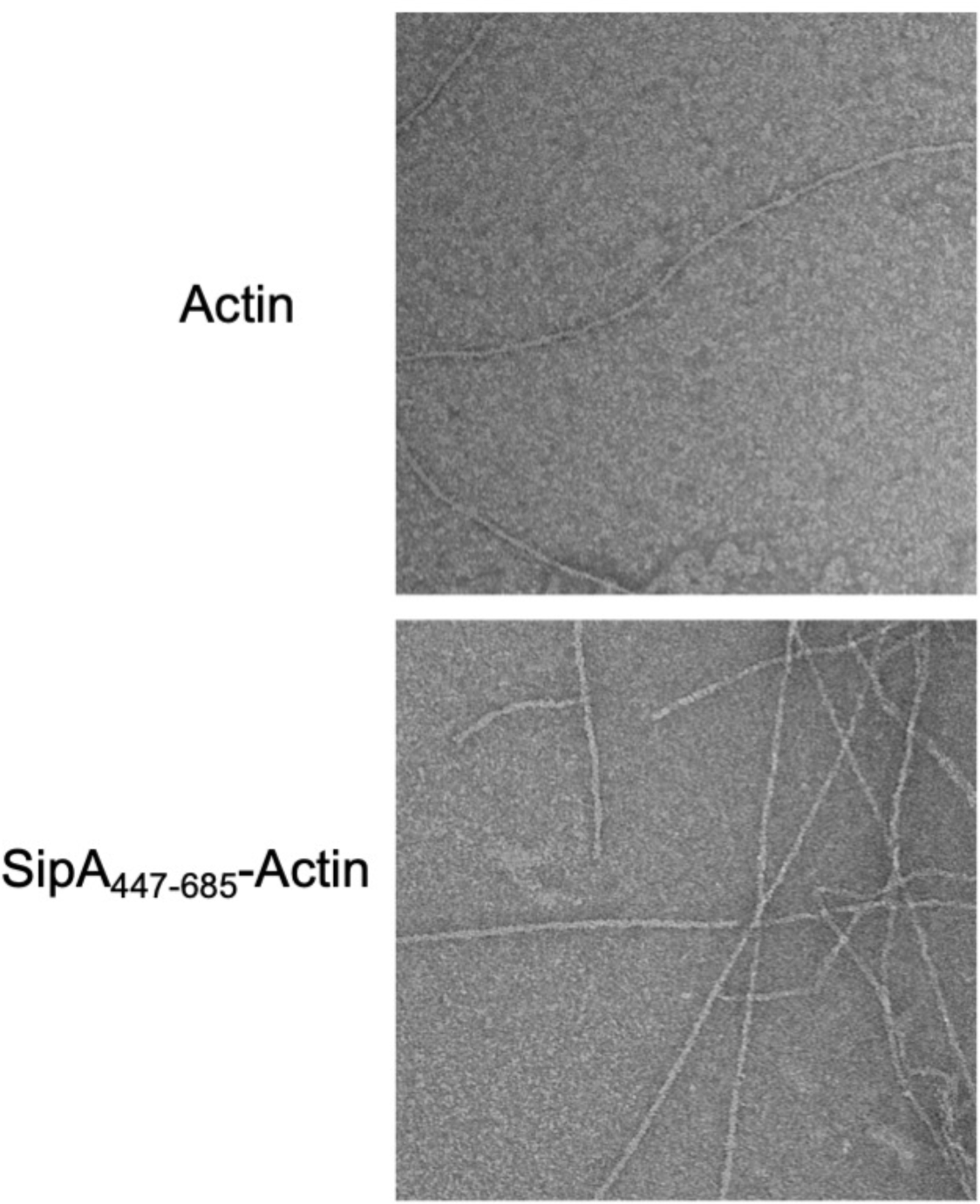
Representative electron micrographs of negatively stained actin filaments alone and with SipA447-685.

**Supplementary Fig. 4.**
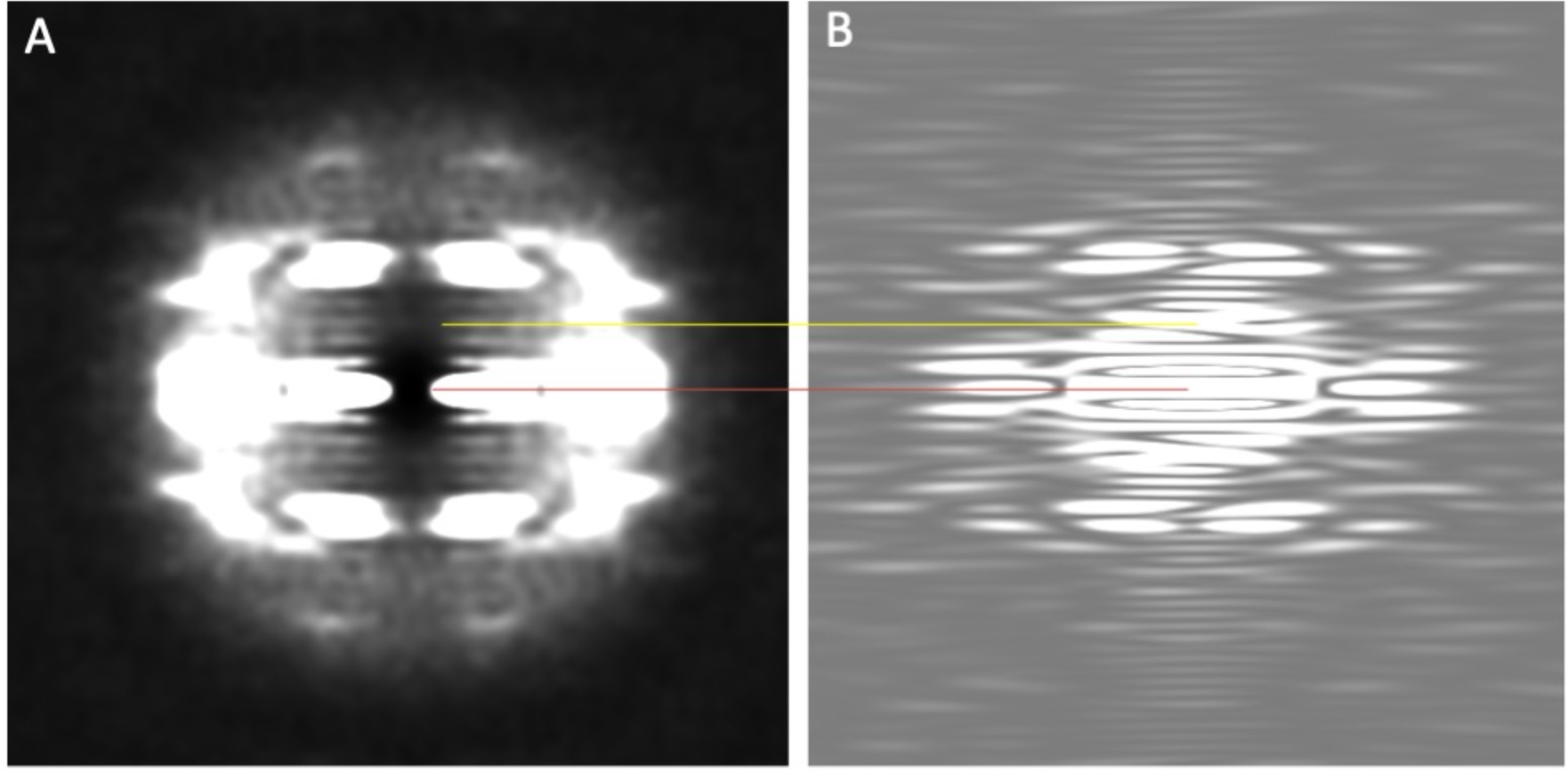
Layer line images of actin filaments decorated with SipA. (**A**) Averaged power spectrum of helix segments boxed out from 1300 micrographs. (**B**) Power spectrum calculated from map projections. The Equator line (n = 0) and the first rise line (n = 0) are indicated with red and yellow lines, respectively. Rise = (pixel size) × (layer-line image size)/ (layer-line height) = 1.068 × 4,096/39.91 = 109.6 Å.

**Supplementary Fig. 5.**
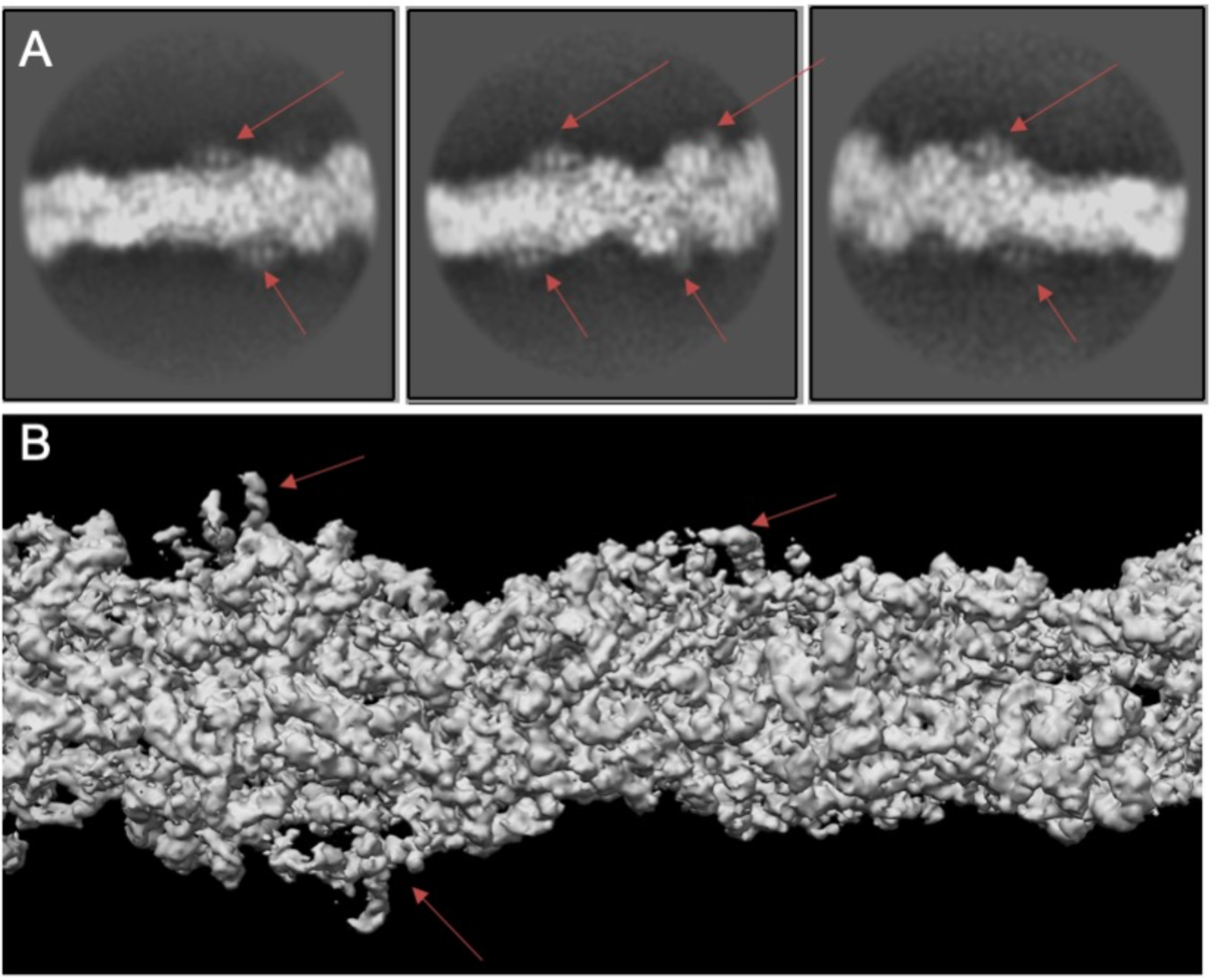
Lower electron map occupancy of SipA_446-684_ in actin filaments. (**A**) representative 2D class averages of SipA_447-685_ decorated actin filaments. Arrows show the density of SipA_447-685_. (**B**) Initial 3D reconstruction of SipA_447-685_ decorated actin filaments. Arrows show the density of SipA_447-685_.

**Supplementary Fig. 6.**
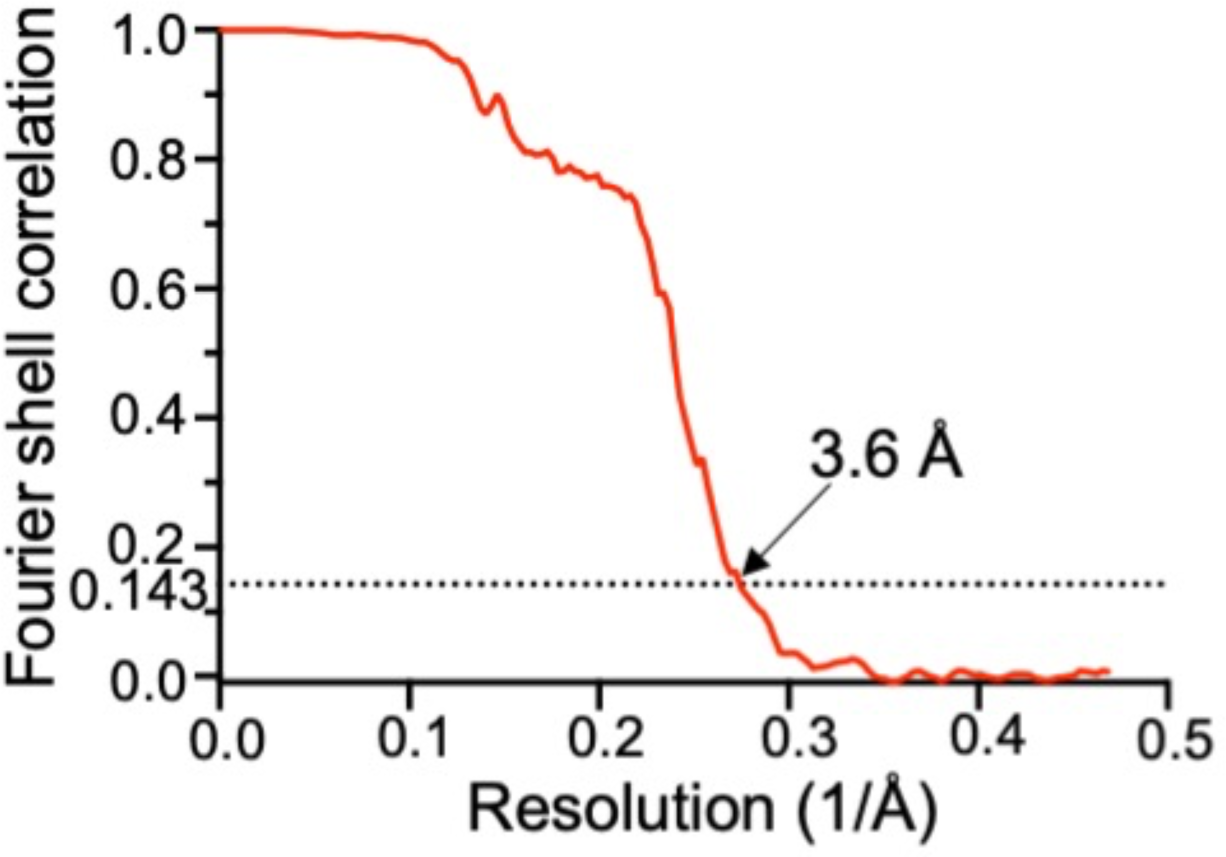
Global resolution estimation of the SipA_446-684_-actin complex. Fourier shell correlation for maps constructed from two half data sets. A cutoff value of 0.143 is used.

**Supplementary Fig. 7.**
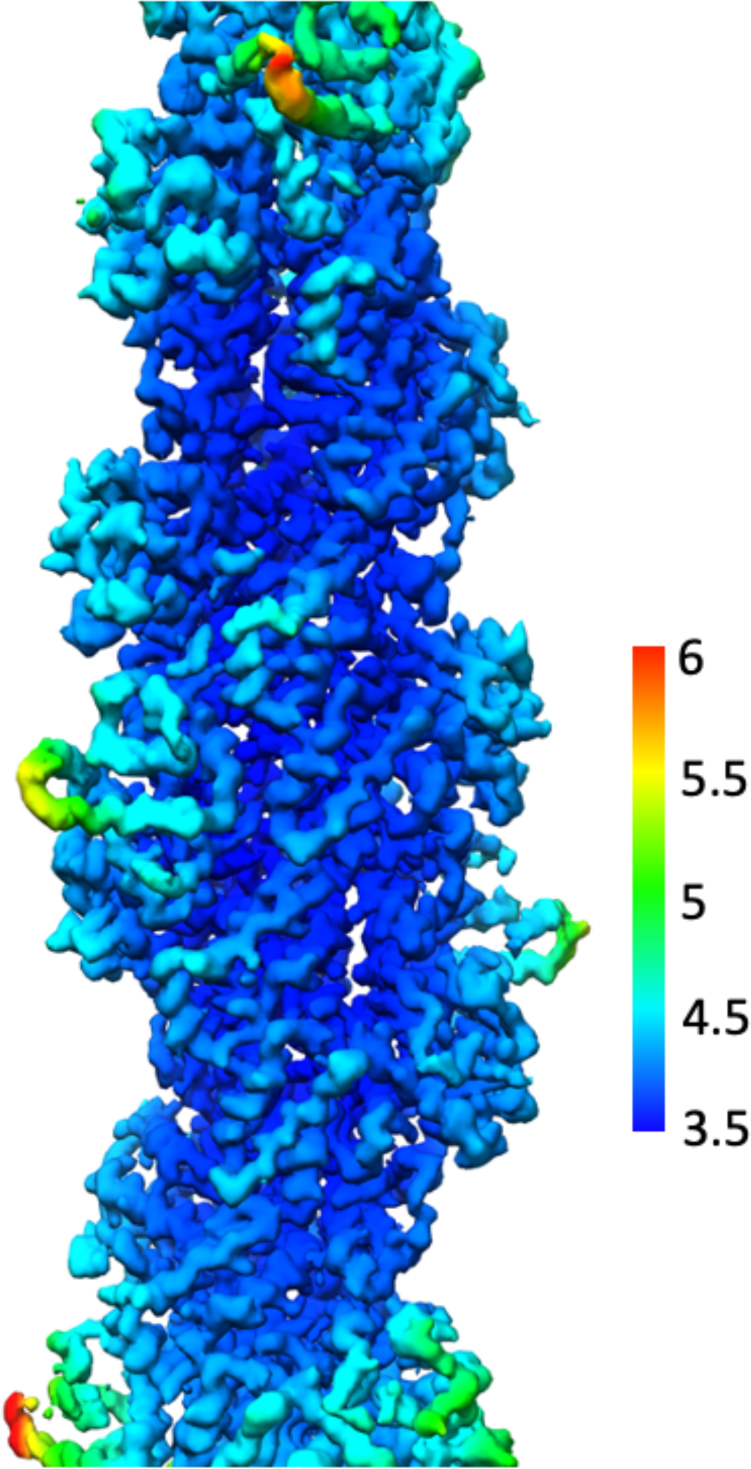
Local resolution estimation for SipA-decorated F-actin with Resmap.

**Supplemental Fig. 8.**
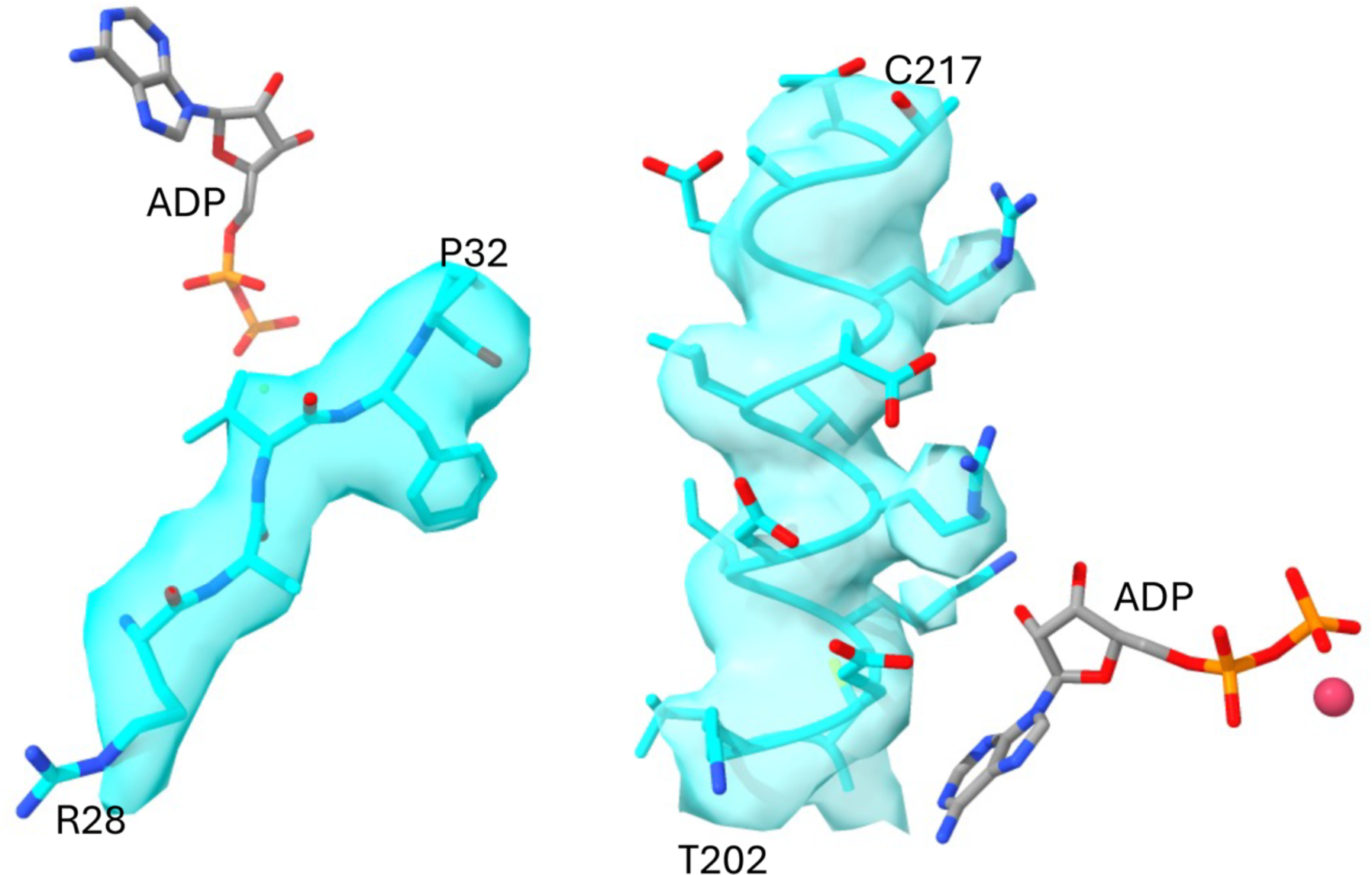
Cryo-EM map and visualization of the side chains of selective residues of actin near the ADP binding site of the model. The density for the fitted coordinates shows features consistent with a resolution of ∼ 3.6 Å.

**Supplemental Fig. 9.**
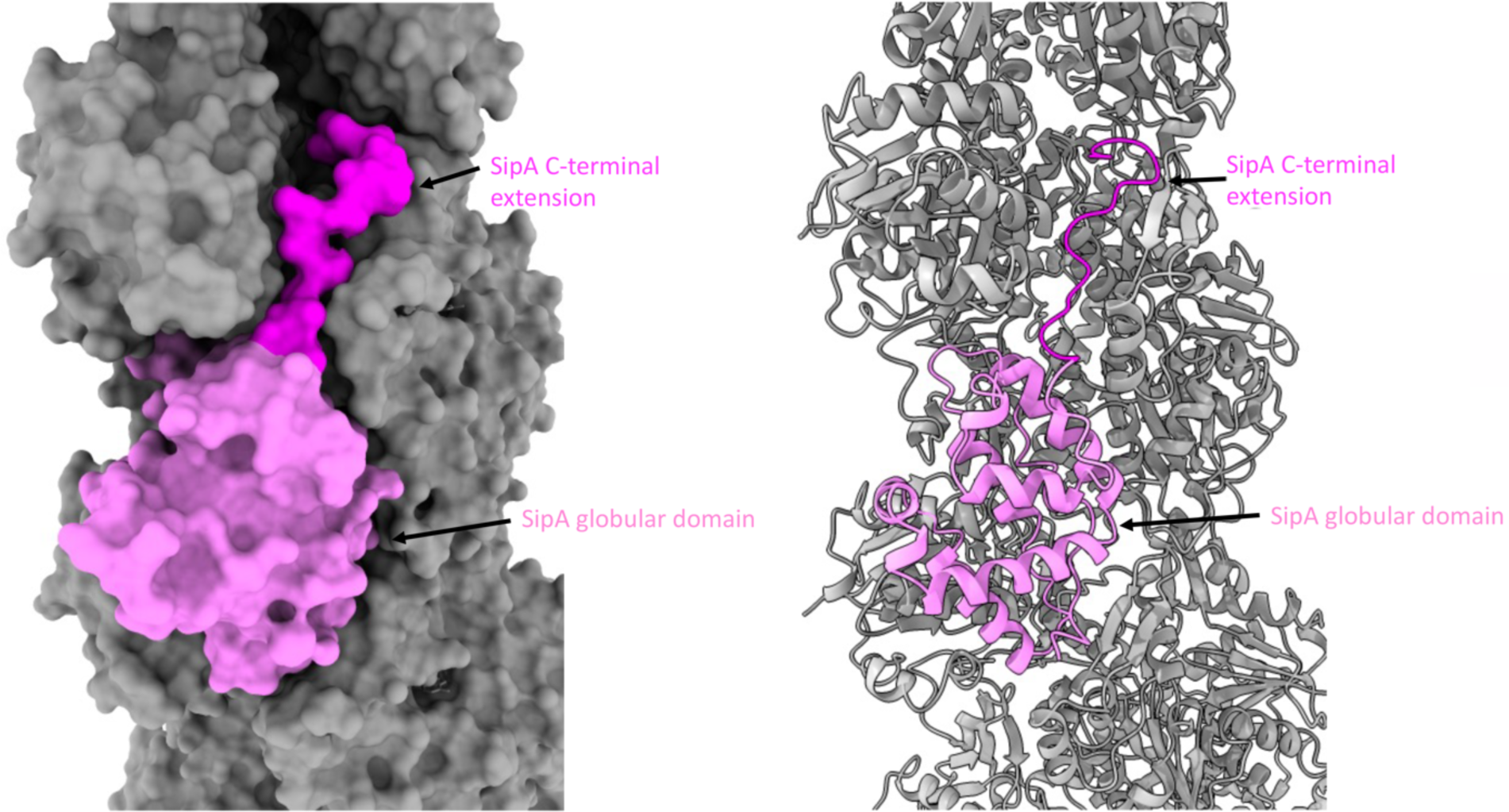
The globular domain (violet) and the C-terminal extension (magenta) of SipA. Actin filament is colored in gray. Ribbon diagram is shown on the right panel.

**Supplementary Fig. 10.**
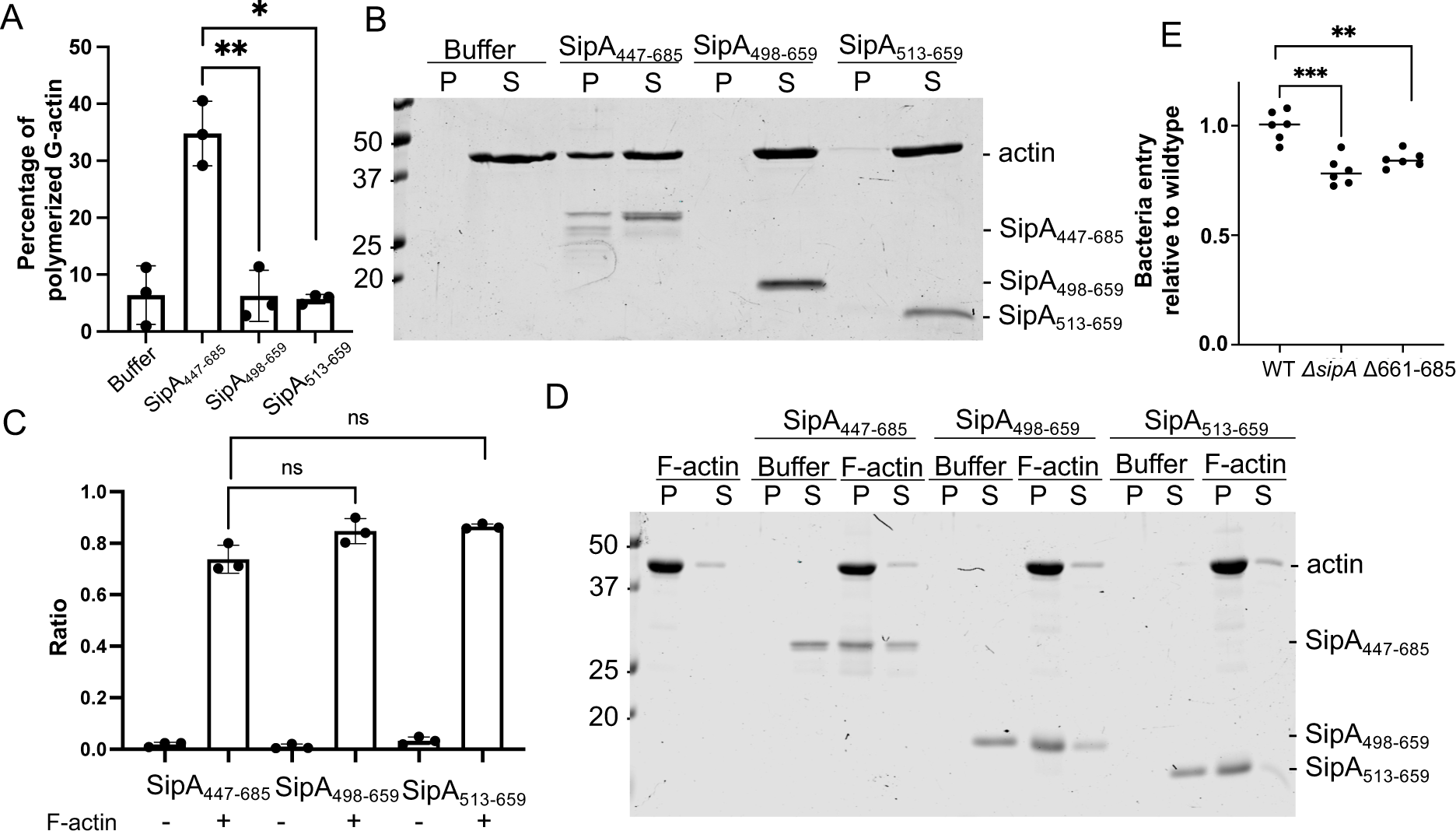
The unstructured arm regions are necessary for the G-actin nucleation function of SipA but are dispensable for its ability to stabilize F-actin. (**A** and **B**) G-actin nucleation activity of different SipA constructs as determined by a co-sedimentation assay. Values shown in (**A**) are the mean ± SD of three independent experiments (*: P<0.05; **: P<0.01; unpaired two-sided t-test). (**C** and **D**) F-actin binding activity of different SipA constructs as determined by a co-sedimentation assay. Values shown in (**C**) are the mean ± SD of three independent experiments (ns: differences not significant, unpaired two-sided t-test). (**E**) A SipA mutant lacking the unstructured C-terminal region exhibits reduced epithelial cell invasion ability. The invasive ability of the different strains was measured by the gentamicin protection assay as described in the methods. Values represent the percentage of the inoculum that survived antibiotic treatment due to internalization and are the mean ± SD of six independent determinations normalized to wild type, which was set to 1. (**: P<0.01; ***: P<0.001; unpaired two-sided t-test). WT: wild type.

**Supplementary Fig. 11.**
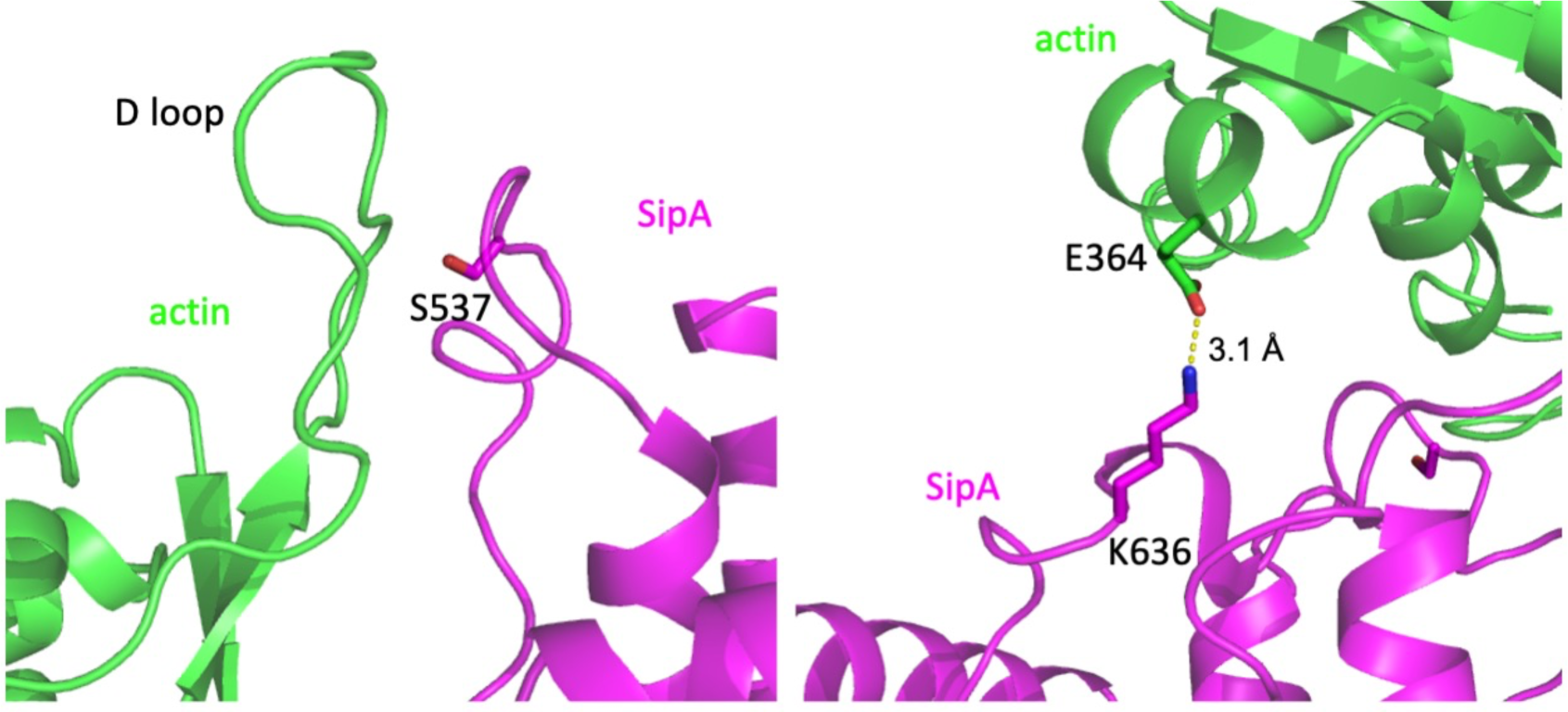
Residues S537 and K636 of SipA interact with the D-loop and C-terminus of actin. Interfaces between SipA (magenta) and actin (green) are shown. Interface residues S537 and K635 of SipA, as well as the D loop and the C-terminal residue E364 of actin are indicated.

**Supplementary Fig. 12.**
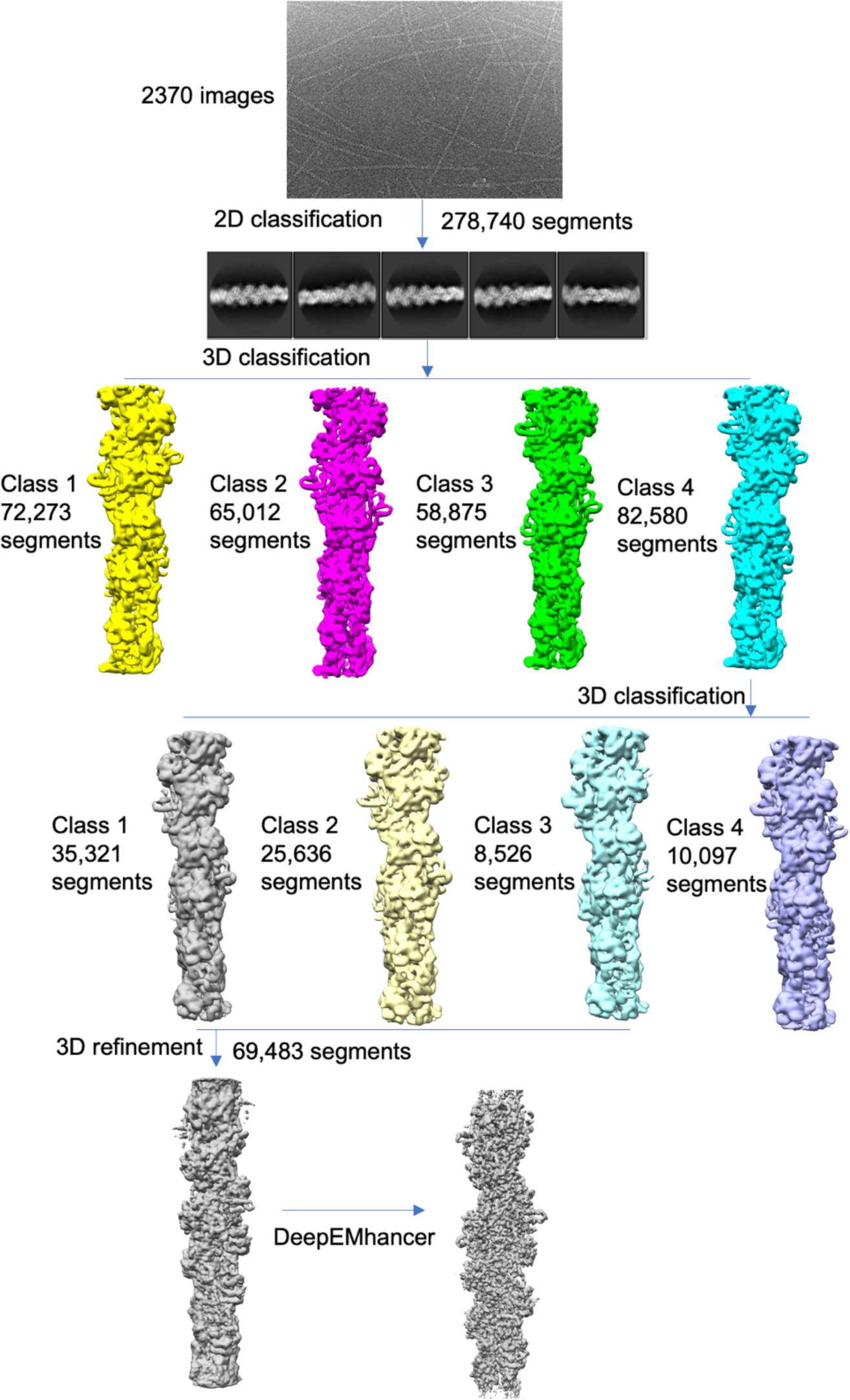
Data processing workflow.

**Supplementary Table S1.**
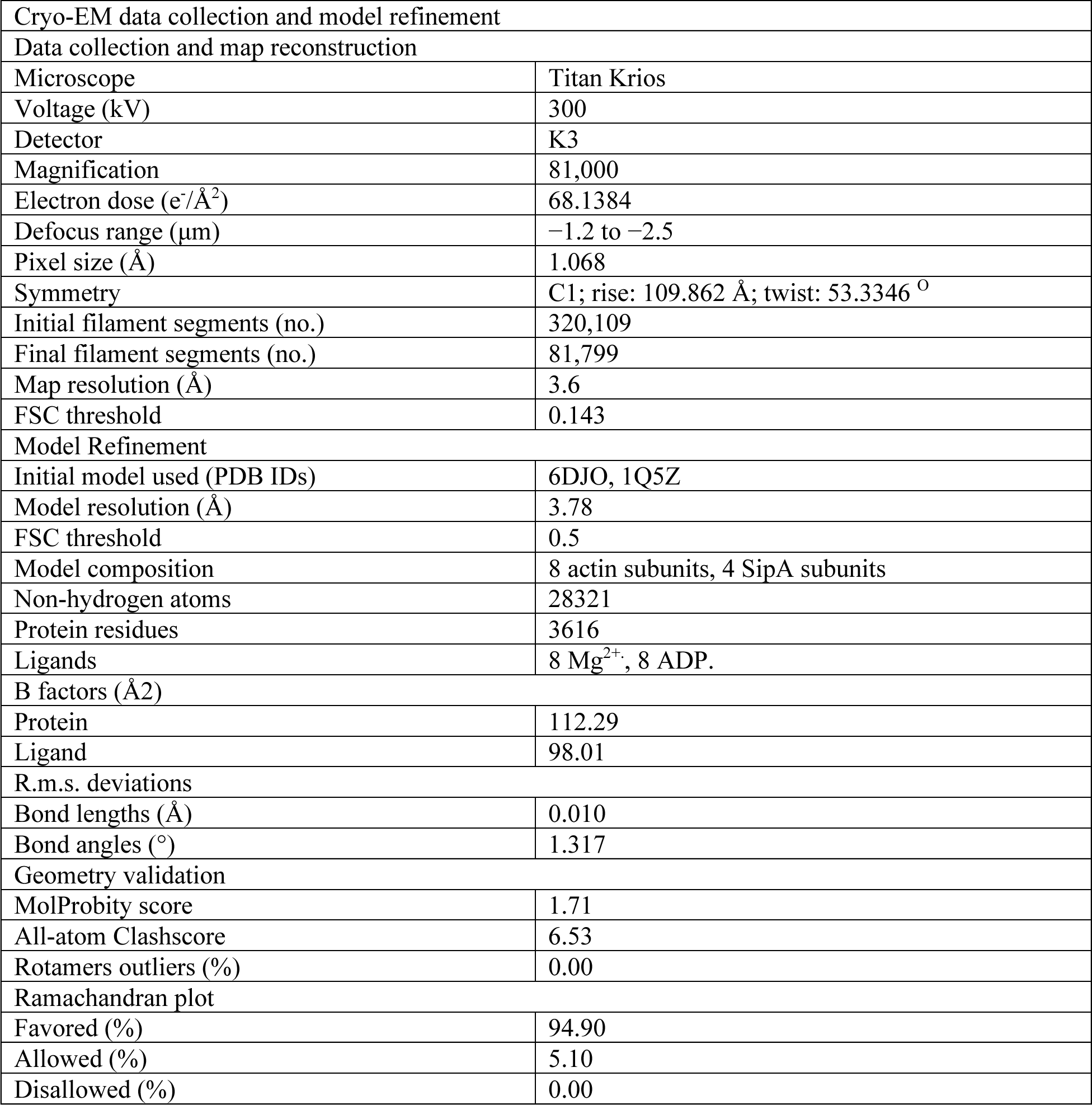
Statistics for data collection, map reconstruction and model refinement.

